# Substrate biasing in UCHL5 proteoforms

**DOI:** 10.64898/2026.07.16.738907

**Authors:** Rishi S. Patel, Nipuni M. Pannala, Chih-Hsuan Lai, Manoj Kumar Sriramoju, Yong-Sheng Wang, Ting Chen, Dabin Jung, Payam Kelich, Soroush Asadi, Elizaveta Shestoperova, Shalini Iyer, Emad Tajkhorshid, Daniel Flaherty, Eric Strieter, Shang-Te Danny Hsu, Chittaranjan Das

**Author notes:** indicates cocorresponding authors To whom all correspondence should be directed. indicates equal contribution (Rishi S. Patel, Nipuni M. Pannala).

## Abstract

Proteolytic deubiquitinating enzymes bridge a gap in substrate recognition through complex regulatory mechanisms. A growing portion of these are accomplished through proteoforms that uniquely control association and diverse sets of cleavage capabilities that relay distinct physiological outcomes. This study describes substrate biasing governed by UCHL5 proteoforms. It demonstrates that N-terminal ubiquitination activates the enzyme towards monoubiquitin substrates, a feature that is conserved across UCHL5 homologs. Crystallographic and spectroscopic data suggest that the N-terminal ubiquitin binds intramolecularly in an allosteric binding site and inhibits branched chain substrate cleavage. Association with Rpn13/Adrm1 relieves this inhibition and reestablishes its ability to debranch, potentially controlling nonspecific debranching compared to retention of needed activity on the 26S proteasome. Collectively, this study describes the molecular basis for substrate selectivity in a deubiquitinating enzyme, an unexplored area in the enzymes that counteract ubiquitin E3 ligases.

## INTRODUCTION

Deubiquitinating enzymes (Dubs) are a class of cysteine and metalloproteases that reverse protein post-translational modification (PTM) by ubiquitin (Ub)^1^. Dub dysfunction is accompanied by many pathologies including a range of cancers^2^, neurological disorders, metabolic and autoimmune diseases^3,4^. The varied functions of the nearly hundred Dubs are well recognized in mediating important physiological outcomes elicited by target deubiquitination^5–9^. The role of Dubs is notably recognized for maintaining a balance between protein degradation and stability. Remarkably, even bacterial and viral pathogens cargo enzymes involved in manipulating the host ubiquitinated proteome in their pursuit of survival, including Dubs from *Legionella*^10–12^, *Chlamydia*^13^ and other pathogens^14–19^ that underscore the importance of the Ub system in the regulation of disease.

These proteases impact nearly all Ub-regulated processes by counteracting the coordinated effort of the three enzymes (Ub activating E1 enzymes, Ub conjugating E2 enzymes, and Ub E3 ligases) that together facilitate ubiquitination of substrate lysine residues^20^. Polyubiquitination gives rise to eight different linkage-specific polyUb chains. Adding to its versatility in structure and function, homotypic and heterotypic polyUb chains come in both linear and branched architectures^21^ and establish widely different proteoform modifications. While proteoform dependencies are also widely observed with E3 ligase maturation and substrate degron recognition^22–27^, this extends to Dubs too. Phosphorylation incorporated Ub dimer substrates of six different linkage types have previously demonstrated discrepancies in Dub activity^28^ as has recently identified lysine carbamylation in Met1 chains^29^. In contrast to substrate proteoforms, previous examples like that of the phosphorylation of A20 by IκKβ conferring a change in linkage specificity resulting in Lys63 chain preferential cleavage over the cognate Lys48 linkage type depict intricate control of PTM guided Dub activity^30^. Proteoforms of Dubs can also take shape in large protein complexes, as seen with pseudoDub maturation in the BRISC and ABRISC complexes^31,32^. Taken together, Dubs achieve a complicated series of functional needs through their proteoforms by PTMs and complex protein assemblies.

While generally dozens of E2 enzymes and hundreds of E3 ligases work coherently, several reports have recently shown that some E2 enzymes function independently of E3 ligases. Apart from the two dedicated hybrid E2/E3 enzymes^33–35^, UBE2W displays E3-independent activity by α-amine methionine ubiquitination^36–38^. Enzymatic control by the aminoacylation of substrate lysine residues is a target for rapid degradation catalyzed by UBE2W^39,40^, highly reminiscent of other processes regulating proteostasis^24,41–44^. Recently, Gly-Gly-Met-specific antibodies have revealed that two substrates for UBE2W happen to be UCHL1 and UCHL5 (or UCH37), members of the Ubiquitin C-terminal Hydrolase (UCH) family of Dubs whose activities are regulated by N-terminal ubiquitination and demonstrate that standalone UBE2W-mediated ubiquitination confers both degradative and nondegradative roles^45^.

Regulation and control of Dub catalytic activity have been a reemerging and widespread theme in recent years^46–49^. This is seen in the UCH family of Dubs as well with neomorphic disease mutations and PTMs including ubiquitination in UCHL1^50,51^ to natural product elicited activation^52^ of UCHL3 and its inhibition by non-catalytic engagement with Lys27-linked diUb^53^. Distinct from these two single domains UCH Dubs, UCHL5 and BAP1 are multidomain UCH enzymes. They carry a catalytic N-terminal UCH domain and helical extension termed the UCHL5-like domain (ULD) that specifically regulate associations to deubiquitinase adaptor domains (DEUBAD) in multi-subunit protein complexes. While the BAP1 ULD is specific for association to the ASXL DEUBAD and its paralogs in the the Polycomb Repressive Deubiquitinase (PR-Dub) complex^54^, DEUBADs found in the 26S proteasomal subunit Rpn13 (or ADRM1) and the INO80 chromatin remodeler complex subunit NFRKB^55,56^ selectively bind UCHL5 ULD^57,58^. Distinct from its canonical, catalytic Ub-binding site, Strieter and colleagues have since discovered a previously uncharacterized Ub-binding site on UCHL5 that is essential for proteasome-associated Ub chain debranching activity at Lys48 branchpoints and its relevance towards proteostasis within cells^59–63^. Despite a broad substrate scope including the Lys48-, Lys6-, Lys11-, and Lys63/Lys48-branched substrates, mutations in α5 and α6 of this noncanonical site reconstituted on the proteasome abolishes degradation of substrates modified with branched Ub chain architectures, emphasizing that debranching is instrumental for substrate degradation.

Loss of UCHL5 and its inhibition overcoming Bortezomib-resistance in multiple myeloma further emphasizes its role in epigenetics^64^ and proteostasis^65^. Unlike the well characterized small molecule inhibitors for UCHL1^66–69^, the only reported UCHL5 inhibitor, bAP-15, is mechanistically ambiguous targeting both cysteine Dubs associated with the proteasome^70^. To address this gap, Strieter and colleagues have recently developed two distinct nanobody inhibitors of UCHL5 that target the canonical and noncanonical sites that inhibit cleavage of monoUb C-terminal substrate and at Lys48 branch points^71^.

Here, we discover a natural means of UCHL5 inhibition that imposes proteoform specific substrate biasing. A UBE2W-mediated N-terminal Ub modification inhibits the ability of UCHL5 to debranch at Lys48 branch points yet activates the enzyme towards monoUb substrates, a feature that implies a proteoform specific role. This deviation in the reaction path further reinforces and supports control of a UCH enzyme, universally believed to process short Ub peptidyl substrates on the account of a sterically limiting crossover loop, in free form as compared to when associated with larger protein complexes. Findings here are consistent with the intramolecular association of the N-terminal Ub with the noncanonical Ub-binding site at α5 and α6 previously demonstrated to be a key recognition area for the Lys48 branchpoint in branched Ub oligomers^60^. We further demonstrate that association with Rpn13 relieves this inhibition and reconstitutes the ability for the N-terminal ubiquitinated proteoform to debranch, effectively suggesting debranching to be retained on the proteasome. This study demonstrates a previously undocumented regulation mode in substrate biasing of a Dub that is noncanonically ubiquitinated by an E2 enzyme and is subject to rules further imposed by its association to large protein complexes.

## RESULTS

### UCHL5 is distinctly activated by N-terminal ubiquitination that is dependent on back-site association

Towards understanding the activation mechanism by N-terminal ubiquitination, profluorogenic minimal substrate cleavage of Ub 7-amino-4-methylcoumarin (Ub-AMC) or Ub rhodamine 110 (Ub-Rho) was used to assess proteoforms of UCHL5 and their mutants. Michaelis-Menten kinetics was determined for UCHL5 WT, Ub^G76V^UCHL5 (Ub-UCHL5), their respective Rpn13^DEU^ complexes, the catalytic domain of UCHL5 (UCHL5^1-228^ from here on denoted as UCHL5 CD) and Ub^G76^UCHL5^1-228^ (Ub-UCHL5 CD) using Ub-AMC (Figure 1A, 1B). While the G76V mutation protects Ub-UCHL5 from self-hydrolysis in vitro, these experiments give a holistic understanding of the modification in the context of UCHL5 proteoforms. The Michaelis-Menten kinetics reported here for UCHL5 CD, UCHL5 and UCHL5 in complex with Rpn13^DEU^ agree with previously reported studies^58,72^ (Figure 1C). In the simplest case, UCHL5 CD and Ub-UCHL5 CD have catalytic efficiencies of 1.61 μM^-^ s^-^ and 21.41 μM^-^ s^-^, suggesting the catalytic domain alone is sufficient for activation by N-terminal ubiquitination and is dependent on both an increase in *k*_cat_ or decrease in K_m_. Activation extends to the full-length enzyme and is comparable to activation imparted by association to Rpn13^DEU^ as they confer nearly a 7-fold and 4.5-fold increase in *k*_cat_/K_m_ respectively. Activation towards other minimal monoubiquitin substrates such as Ub-Rho is also observed (Supplementary Figure 1A, 1B), perhaps suggesting a versatility in engaging varied monoUb C-terminal conjugates. With a catalytic efficiency of 9.98 μM^-^ s^-^, UCHL5 is activated more by N-terminal ubiquitination than association with Rpn13^DEU^ (*k_c_*_at_/K_m_ of 6.22 μM^-^ s^-^) towards Ub-AMC hydrolysis despite a similar K_m_. Inactivation kinetics using the suicide inhibitor Ub glycine vinylmethylester (Ub-VME) suggest that at low concentration ([S] < K_m_) of substrate, both Ub-UCHL5 and UCHL5 Rpn13^DEU^ exhibit similar reactivity with a *k*_inact_/K_i_ of 0.38 μM^-^ s^-^ and 0.36 μM^-^ s^-^ (Supplementary Figure 1C). Notably, these two activation mechanisms are synergistic. The Ub-UCHL5 Rpn13^DEU^ complex has the highest catalytic efficiency of the enzymes tested at 23.65 μM^-^ s^-^. A lower *k*_cat_ is observed with the Ub-UCHL5 Rpn13^DEU^ complex, indicating that activation is driven more by K_m_ (820 nM) rather than the *k*_cat_ effect as seen in the case of Ub-UCHL5 and UCHL5 Rpn13^DEU^ individually.

**Figure 1.**
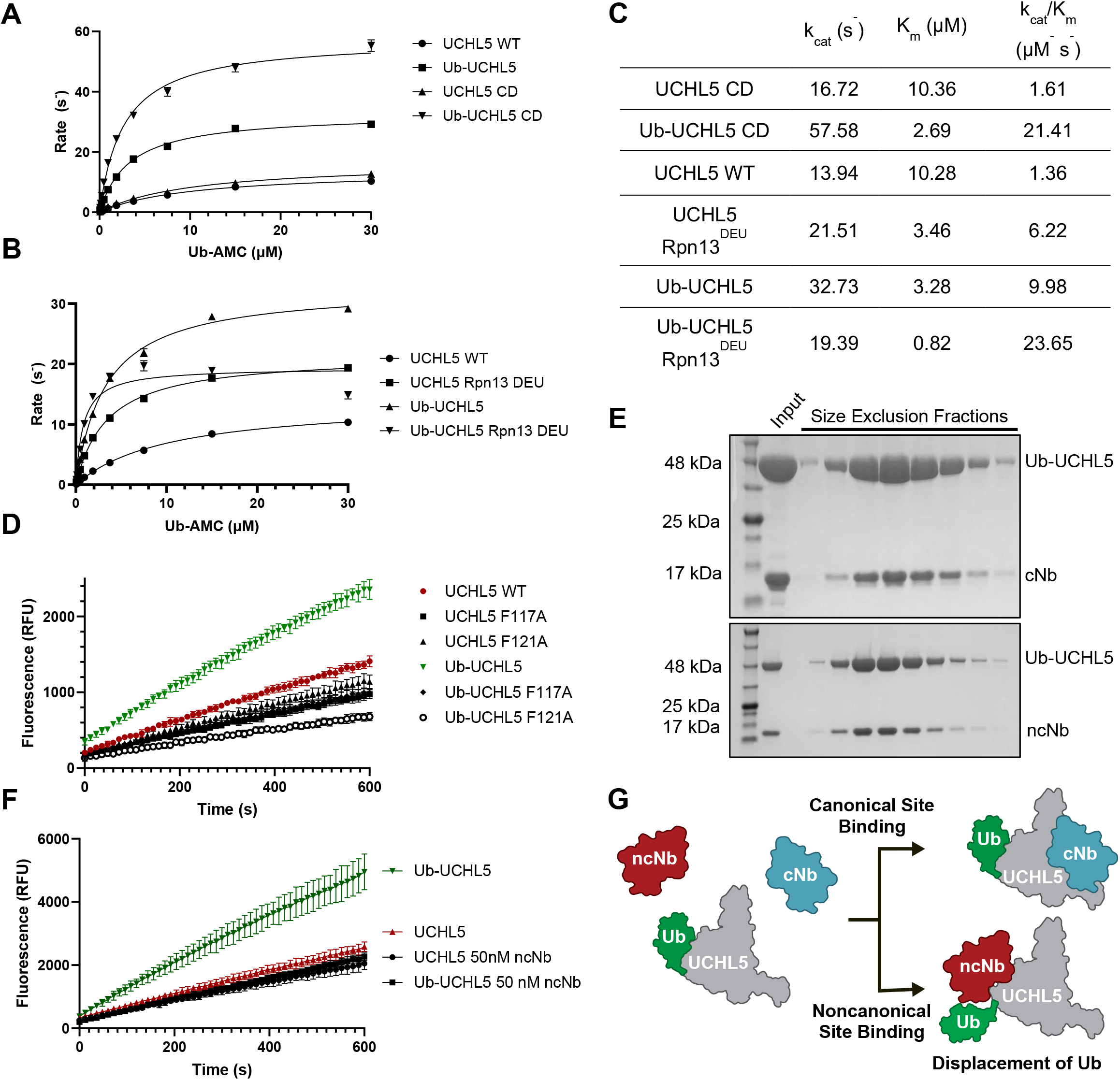
Activation of UCHL5 by N-terminal ubiquitination is dependent on an allosteric Ub binding site. Michaelis Menten kinetic plots for the catalytic domain and full length UCHL5 and N-terminally ubiquitinated UCHL5 indicating activation is independent of the ULD (A). Michaelis Menten kinetic plot comparing activation by N-terminal ubiquitination to the association of Rpn13^DEU^ and indicating a synergistic activation of both modes (B). (C) A table summarizing the Michaelis Menten kinetic parameters for enzymes tested in (A) and (B). Single concentration Ub rhodamine 110 assay against mutant UCHL5 and Ub-UCHL5 reveals two key phenylalanines F117 and F121 in the allosteric Ub binding site are important for activation (D). SDS-PAGE Coomassie-stained gel of size exclusion chromatography fractions show coelution of both nanobodies that bind in the canonical and noncanonical sites (E). Single concentration Ub rhodamine 110 assay illustrating a nanobody that binds in the allosteric Ub binding site inhibits activation imparted by N-terminal ubiquitination (F). Schematic representing nanobody binding to Ub-UCHL5, particularly the displacement of intramolecular association and its inhibitory effect on activation is shown in (G).

Structural comparisons of previously solved UCHL5 structures available in the Protein Data Bank depict the N-terminus extending towards the region termed as the “back-site” or noncanonical Ub-binding site^73^ (Supplementary Figure 3B). We reasoned that intramolecular association of N-terminally modified Ub occupies the same noncanonical Ub binding site presented by α5 and α6 of UCHL5. With this site bearing two key residues, F117 and F121, mutations to alanine have previously been shown to abolish debranching activity in vitro and reconstituted on the 26S proteasome^74^. These same mutations, F117A and F121A, impair activation by N-terminal ubiquitination towards Ub-Rho, restoring the activity to levels observed with either UCHL5 WT or the corresponding mutant in the unmodified background (Figure 1D). This data suggests that, instead of projecting away from the UCH domain, the N-terminal Ub inadvertently occupies the noncanonical site through intramolecular folding, aided by both the proximity of the N-terminus of UCHL5 to the site and the flexibility afforded by the C-terminal tail of Ub. Thus, the previously known noncanonical site also serves as an allosteric Ub binding site.

Further investigations using the two nanobodies that bind at either the canonical site (cNb) or the noncanonical site (ncNb)^71^ were carried out by determining coelution of the incubated protein mixtures by size exclusion chromatography. We hypothesized that occupation of the noncanonical site by the N-terminal Ub would prevent the association of ncNb with Ub-UCHL5 while still maintaining affinity for cNb. However, both the cNb and ncNb, whose affinities are 200 nM and 150 pM respectively, coeluted with Ub-UCHL5 (Figure 1E). Demonstrated by Ub-UCHL5’s ability to readily react with the S1 site directed Ub-VME probe, the cNb is expected to bind to UCHL5 since the proximally located S1 site remains unperturbed. However, the unusual high affinity for ncNb towards UCHL5 and its coelution with the ubiquitinated form warranted further investigation. Previously reported hydrogen-deuterium exchange experiments have found that the ncNb binds in the regions described by residues 114-135, residues that comprise the cryptic binding site^71^ (including residues F117 and F121). Surprisingly, treatment of ncNb with Ub-UCHL5 in the Ub-Rho cleavage assay impaired activation, lowering the activity to levels observed with unmodified UCHL5 (Figure 1F). This suggests that ncNb outcompetes the intramolecular association of the N-terminal Ub (Figure 1G), while indicating that the activation is mediated by a Ub-specific mechanism that is consistent with binding in the cryptic Ub-binding site.

To determine the molecular basis of N-terminal Ub-mediated regulation of UCHL5, we solved the X-ray crystal structure of Ub-UCHL5 to a resolution of 2.7 Å in the orthorhombic space group C222_1_. Four chains of Ub-UCHL5 are observed in the asymmetric unit. All four chains reveal Ub bound to the backside of the protein (Figure 2A) that lies opposite the canonical S1 binding site of UCHL5 (Figure 2B). To our knowledge, this represents the first structural characterization of Ub engaging the backside of UCHL5, uncovering a mechanism by which it remodels substrate specificity. Rather than projecting freely from the N terminus, the Ub moiety folds back onto the UCHL5 surface, forming a well-ordered intramolecular interface that buries approximately 890 Å^2^ of solvent-accessible surface area (SASA). The Ile44 hydrophobic patch of the N-terminal Ub engages the helix-loop-helix motif of α5 and α6 on UCHL5 through surface-exposed hydrophobic residues Phe117 and Phe121 packing against Ile44 and Leu8 of the Ile44 hydrophobic patch on Ub (Figure 2C). Additional hydrophobic contacts are observed between Met125 and Leu128 on UCHL5 with Ile44 and Val70 on Ub. Observed at the interface are three direct hydrogen bonds (Figure 2D), observed between backbone amide of Ub Leu73 and the side chain of UCHL5 Asn132, side chain of Ub His68 and the backbone carbonyl of UCHL5 Phe117, and the backbone carbonyl of Ub Gly47 and the backbone amide of UCHL5 Asp122. In this structure, two of the four chains show a pronounced deviation of the C-terminal tail, adopting an approximately 180° reoriented conformation that is likely attributable to crystal packing effects. In three of the four UCHL5 protomers, Cys88 lies within hydrogen bonding distance of His164. This is consistent with a catalytically competent active site observed in apo and substrate-bound UCHL5 structures (Supplementary Figure 3D). However, the fourth protomer adopts a misaligned conformation that is consistent with active site geometries seen with UCHL1 catalytic triad misalignment^75,76^, suggesting that modified UCHL5 samples multiple conformational states in solution (Figure 2E). While it is not clear how this relates to activation, this also represents the first crystallographic depiction of proper catalytic triad misalignment observed in UCHL5. Finally, it was observed that electron density of residues in the crossover loop (155-161) and the linkage between Ub and Met1 of UCHL5 are both poorly defined, consistent with intrinsic flexibility in these regions.

**Figure 2.**
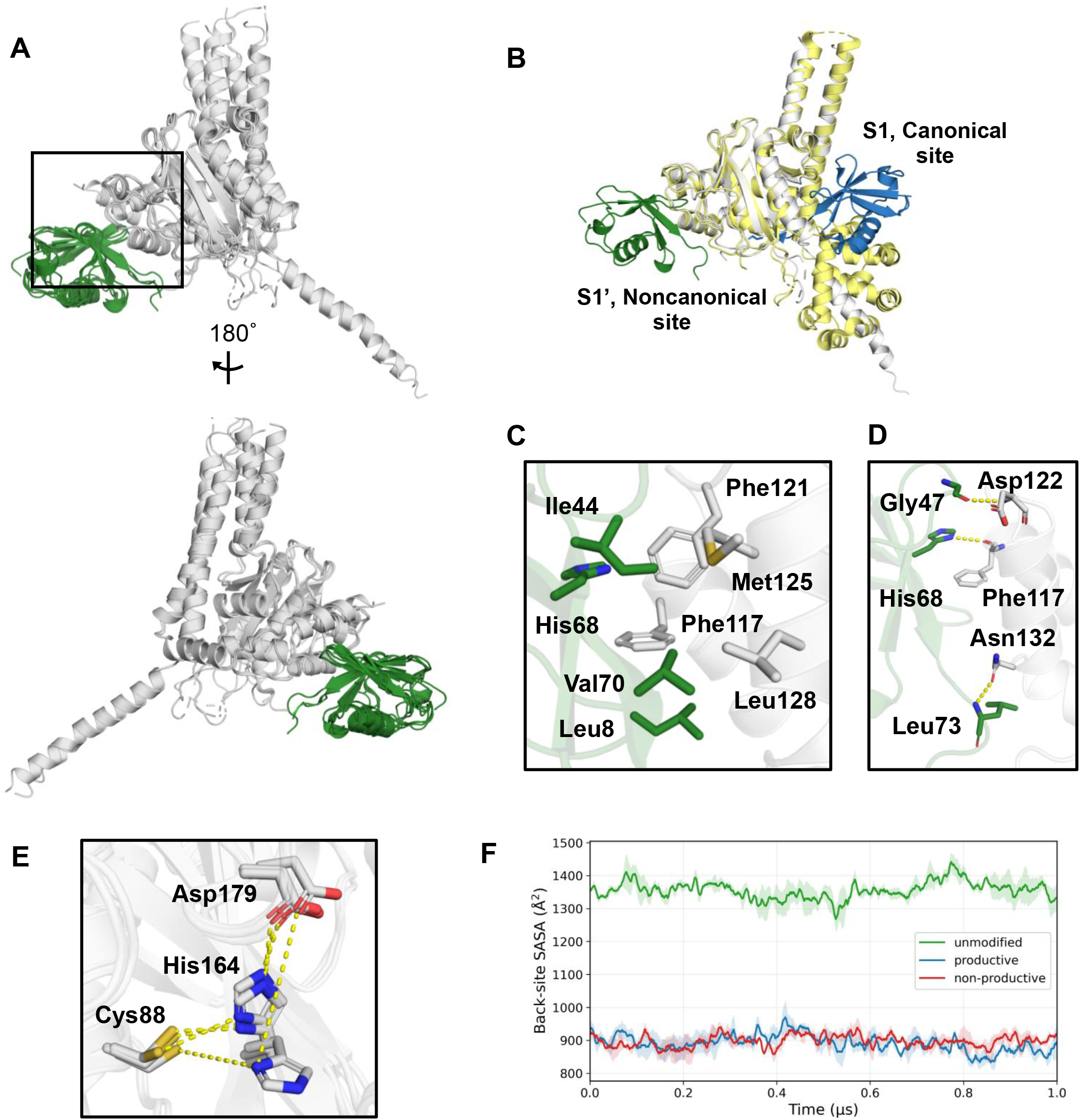
X-ray crystallography and molecular dynamics simulations of Ub-UCHL5 describe exposed hydrophobic regions responsible for mediating interactions. Crystal structure of Ub-UCHL5 solved to a resolution of 2.7 Å. The four chains of Ub-UCHL5 found in the asymmetric unit super imposed onto one another depicts Ub binding in the allosteric Ub binding site (A). The N-terminal Ub is found to take a different trajectory in the comparative overlay of an individual protomer found in the asymmetric unit of the solved crystal structure onto the structure of Ub-bound UCHL5 Rpn13^DEU^ structure (B). The intramolecular interaction between the N-terminal Ub and UCHL5 is driven by hydrophobic interactions occurring between the Ile44 hydrophobic patch on Ub and hydrophobic residues F117, F121, Met125 and Leu128 on UCHL5 (C). There are three intramolecular hydrogen bonding interaction that further stabilize this interaction (D). The active site catalytic triad in C88, H164, and D179 across all the chains found the asymmetric unit show a relative aligned conformation (E). Back-site solvent-accessible surface area (SASA) versus time over the first 1 µs molecular dynamics simulations (F) with mean ± SEM over three replicates.

To determine whether the crystallographic N-terminal Ub orientation was natural or a product of crystal contact, we performed all-atom molecular dynamics (MD) simulations of the systems in explicit water to glean atomic insights into the effect of ubiquitination. Beginning from the unmodified enzyme and the two distinct active-site conformations of Ub-UCHL5 that include an aligned, active-like (productive) protomer and a misaligned (non-productive) protomer, we ran three independent 1-µs simulations for each system. Across all trajectories the UCHL5 core remained folded and stable (Cα RMSD of approximately 3-4 Å with modest replica-dependent excursions for all three systems), and the appended N-terminal Ub remained docked at the noncanonical back-site throughout the simulations. The Ub Ile44 hydrophobic patch stayed in persistent contact with F117 and F121 of UCHL5 at a near-constant center-of-mass distance of ∼9 Å in both productive and non-productive protomers, and back-site solvent accessibility was reduced by approximately one-third relative to the unmodified enzyme (Figure 2F). These observations corroborate the crystallographic findings that the N-terminal Ub occupying the α5– α6 back-site by its Ile44 patch and show that this engagement is maintained in solution.

The effect of N-terminal ubiquitination on the structure and dynamics of UCHL5 in solution was investigated by multidimensional heteronuclear nuclear magnetic resonance (NMR) spectroscopy. We first examined the two-dimensional (2D) ^15^N-^1^H and ^13^C-^1^H heteronuclear single quantum coherence (HSQC) spectra of uniformly ^13^C and ^15^N labeled, [U-^13^C/^15^N], full-length UCHL5 fused with ubiquitin at the N-terminus. However, the spectrum exhibited severely broadened signals due to the high molecular weight (Supplementary Figure 5A). We, therefore, opted for the UCHL5 CD, and performed segmental stable isotope labeling to introduce ^13^C/^15^N labeling to either Ub or UCHL5 CD followed by sortase-mediated ligation^77,78^ with unlabeled components to minimize the spectral overlap in the fingerprint ^15^N-^1^H HSQC spectra. The ligated product retained activation (Supplementary Figure 5B). Superposition of the 2D ^15^N-^1^H HSQC spectra of ubiquitin before and after ligation with UCHL5 CD and counterpart in UCHL5 CD showed significant spectral changes with many cross peaks exhibiting sizeable chemical shift perturbations (CSPs), indicating changes in the microenvironments around the amide groups of these effected residues (Figure 3A, 3B).

**Figure 3.**
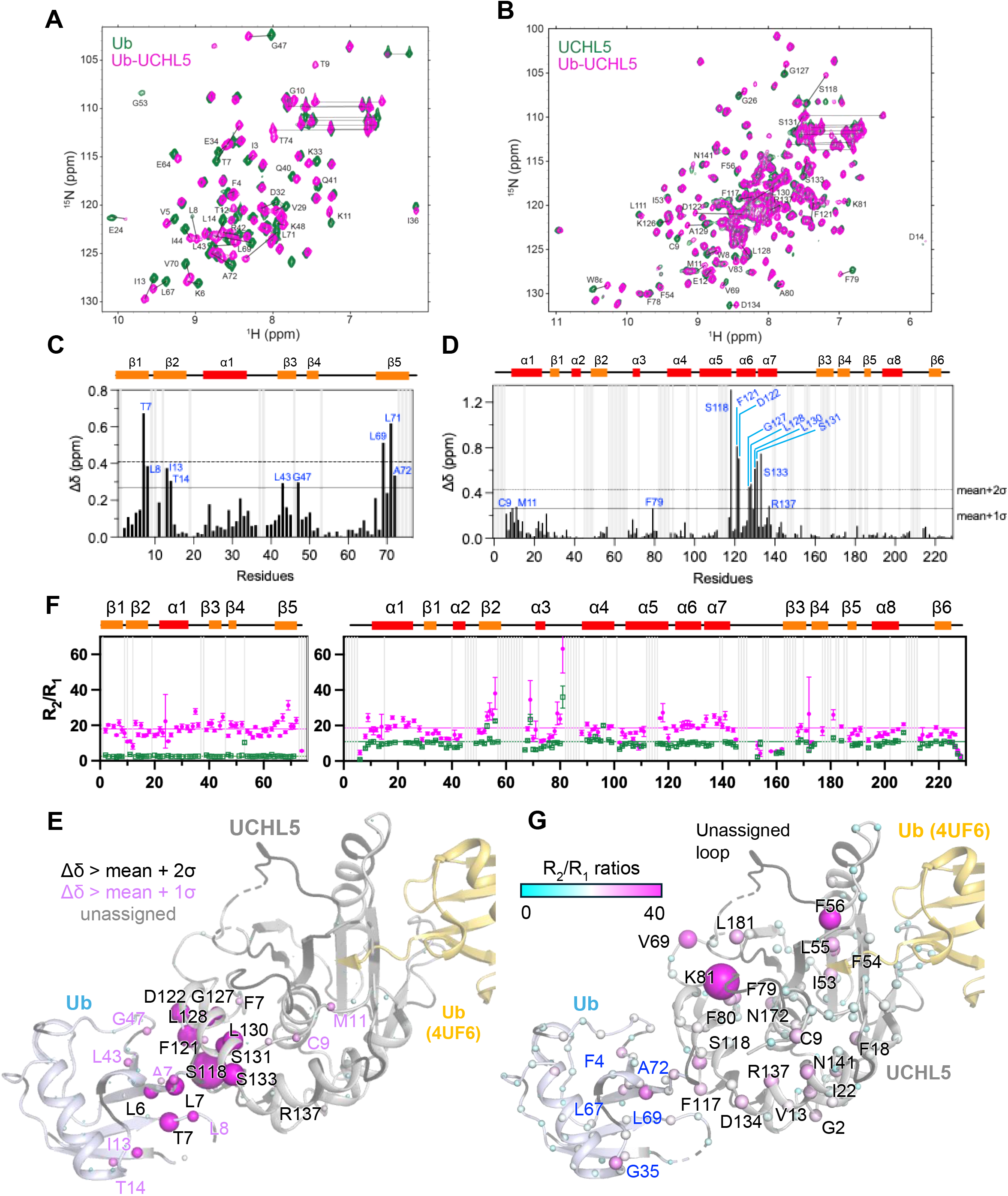
Structural mapping of NMR chemical shift perturbation and 15N relaxation rates in Ub and UCHL5 CD upon sortase-mediated ligation. Superposition of the ^15^N-^1^H HSQC spectra of Ub before (green) and after (magenta) ligation (A). Superposition of the 15N-1H HSQC spectra of UCHL5 CD before (green) and after (magenta) ligation (B). Residues showing significant chemical shift perturbations are indicated with the residue identities. Weighted chemical shift perturbations (Δδ) as a function of residue number in Ub (C) and UCHL5 CD (D). The thresholds for the mean values plus one and two standard deviations are indicated in solid and dashed lines, respectively. The identities of the residues that exhibit significant chemical shift perturbations above the thresholds are indicated. The R_2_/R_1_ ratio of Ub (left) and UCHL5 CD (right) as a function of residue number without and with ligation are shown in open squares and filled magenta circles (E). The average R_2_/R_1_ ratios derived from residues within the secondary structural elements are shown in horizonal lines with matching colors as defined in Supplementary Figure 5B. Structural mapping of the observed chemical shift perturbations (F) and R_2_/R_1_ ratio (G) using the crystal structure of Ub-UCHL5 fusion shown in gray cartoon representation. The previously reported UCHL5-bound Ub structure shown in yellow (PDB ID 4UF6) is superimposed with the structure UCHL5 to illustrate the relative orientation of the bound substrate. The backbone amide nitrogen atoms are shown in sphere with their radii scaled in proportion to the observed chemical shift perturbation (van der Waals radius multiplied by 0.1 of the weighted chemical shift perturbation). Residues that exhibit two standard deviations and one standard deviation above the mean values in Ub and UCHL5 CD are labeled with their identities in black and purple, respectively.

We recorded a set of backbone triple resonance NMR experiments to assign the chemical shifts of backbone amide hydrogen (HN), nitrogen (N), Cα and Cβ atom of Ub and UCHL5 CD before and after ligation. The use of perdeutration in addition to ^13^C/^15^N labeling of UCHL5 CD ([U-^2^H/^13^C/^15^N]-UCHL5 CD) for the ligation with Ub was necessary because of the poor magnetization transfer to the majority of the Cβ atoms in the doubly ^13^C/^15^N labeled UCHL5 CD ligated with Ub due to the high molecular weight. Perdeuteration significantly improved the spectral quality of the triple resonance backbone assignment experiments to enable efficient backbone resonance assignments. The composite backbone amide CSPs were plotted as a function of sequence number of Ub fused with UCHL5 CD (Figure 3C, 3D). The CSPs were the highest for T7, L69 and L71 in Ub and for S118, F121, D122, G127 L128, L130, S131 and S133 in UCHL5 CD, which collectively formed a cluster of residues localized within the interface between the two proteins as observed once mapped onto the crystal structure crystal structure (Figure 3E).

We next carried out ^15^N spin relaxation dynamics experiments—longitudinal relaxation (R_1_) and transverse relaxation (R_2_) experiments—of Ub and UCHL5 CD before and after sortase mediated ligation. Before ligation, Ub and UCHL5 CD each exhibited distinct R_1_ and R_2_ values with fairly homogeneous R_2_/R_1_ ratios across the sequences, except for residues flanking α-helix 3 (residues 66-76) and those residing the cross-over loop (residues 150-160) in UCHL5 CD (Figure 3F and 3G). The auto-correlation times of free Ub and UCHL5 CD were 3.71 ± 0.36 ns and 9.93 ± 1.20 ns, respectively (Material and Methods). Upon ligation, the overall R_2_/R_1_ ratios of Ub and UCHL5 CD were very similar, indicating that the two proteins behaved as a single entity in solution, sharing the same molecular tumbling behavior due to the ligation; indeed, the corresponding auto-correlation times of ligated Ub and UCHL5 CD were 13.02 ± 1.63 ns and 13.28 ± 1.66 ns, respectively (Supplementary Figure 5B). While the overall R_2_/R_1_ ratios are relatively homogeneous for Ub, UCHL5 CD exhibited elevated R_2_/R_1_ ratios in the loop connecting β2 and α3 and the loop connecting α3 and α4 compared to the unligated UCHL5 CD. Furthermore, residues 130-141 also exhibited higher R_2_/R_1_ ratios compared to the unligated counterpart; in contrast, residues 30-43 exhibited lower R_2_/R_1_ ratios compared to the unlighted counterpart. These additional relaxation dynamics changes are indicative of either allosteric dynamics changes or alternative inter-domain contacts that are not captured by the crystal structure.

### Activation by Ub binding in the allosteric site is a convergent feature observed in other UCHL5 homologs

With each member of the UCH family bearing N-terminal UCH domains, we next asked whether this modification affects catalytic activities of the other members as well. In contrast to UCHL5, N-terminal ubiquitination exerts an inhibitory effect on the single domain hydrolases UCHL1 and UCHL3 (Figure 4A, 4B), drastically reducing the *k*_cat_/K_m_ towards the Ub-AMC minimal substrate. While internal lysine and even N-terminal ubiquitination is well known to occur on UCHL1, this agrees with the inhibitory action of UBE2W mediated N-terminal ubiquitination of UCHL1, as previously reported by Koerber and colleagues^45^. However, this is not observed with BAP1, whose UCH domain is closely related to UCHL5 and shares 44% sequence identity. BAP1 exhibits a 3-fold increase in catalytic efficiency upon N-terminal ubiquitination (Figure 4C). With the K_m_ marginally changing, the change in activation observed can be attributed primarily to a *k*_cat_ effect (3.30 s^-1^ to 11.29 s^-1^) (Figure 4D). An allosteric activation mechanism of the PR-Dub complex was recently reported wherein monoubiquitination at the C-terminal periphery of ASXL1 DEUBAD at K351 promotes nucleosomal deubiquitination at H2A K119^79–81^. Surprisingly, when overlayed with the structure of BAP1 bound to monoubiquitinated ASXL1 (PDB 9U5U), Alphafold3 predictions of the N-terminally ubiquitinated BAP1 UCH domain in complex with Ub show that there is a converging mechanism that exploits allosteric Ub binding in the back-side similar to that observed with UCHL5. The structure and the prediction alignment with striking similarity and observe a RMSD of 1.5 Å in Cα alignment (Figure 4E). Homologous to F117 and F121 in UCHL5, both F118 and F122 in BAP1 maintain the hydrophobic interface with the back side Ub. While it is unknown whether BAP1 is N-terminally ubiquitinated, this collectively represents a larger importance of converging mechanisms by allosteric Ub binding elicited by proteoform-relevant enzyme regulation.

**Figure 4.**
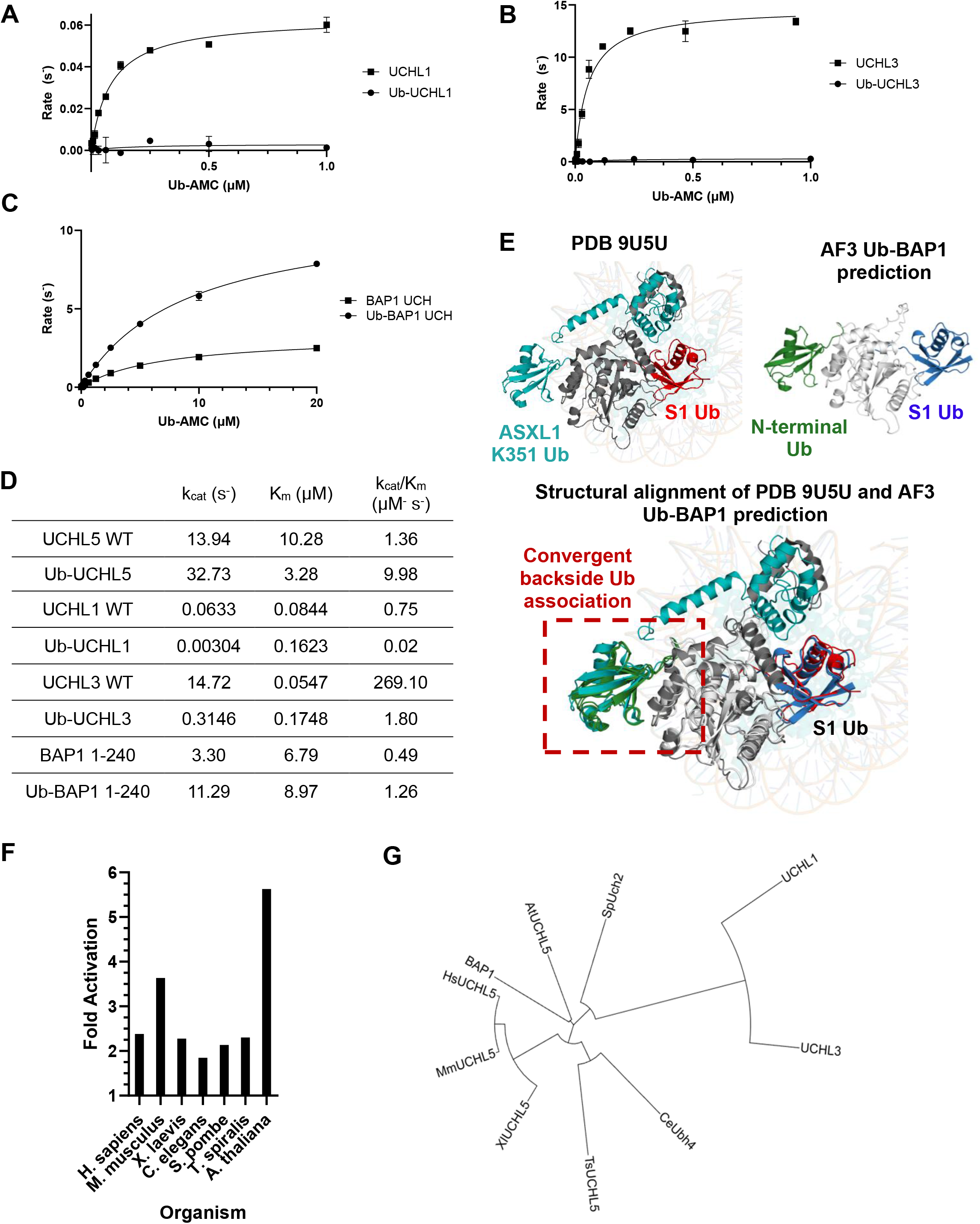
Activation by back side binding is conserved feature in UCHL5 homologs. Michaelis Menten kinetics plots describing the inhibitory effects of N-terminal ubiquitination on UCHL1 and UCHL3 (A, B) yet N-terminal ubiquitination activates the UCH domain of BAP1 (C). (D) A table describing the Michaelis Menten parameters for the enzymes tested in (A, B, and C). Ribbon representation of the cryogenic electron microscopy structure of nucleosome reconstituted BAP1 bound to ubiquitinated ASXL1 on K351 (BAP1: gray/ubiquitinated ASXL1:teal/canonical Ub (red), PDB 9U5U, top left), the AlphaFold3 prediction of N-terminally ubiquitinated BAP1 (noncanonical Ub:green/BAP1:white) bound to Ub (canonical Ub:blue, top right), and their overlay with an alpha carbon RMSD of 1.5 Å (E). Extent of fold activation imparted by N-terminal ubiquitination of UCHL5 homologs from other organisms in single concentration Ub rhodamine 110 assays (F). Phylogenetic tree of UCH domains describes a distinct evolutionary divergence from single domain and multidomain UCH enzymes (G).

Noticing this conservation between two human UCH homologs, we asked whether this feature is evolutionarily conserved. Seemingly, UBE2W is conserved as far back as yeast with UBC16 as its closest ortholog^37^, suggesting that this modification could be an early regulatory mechanism. Homologs of UCHL5 in *M. musculus*, *X. laevis*, *C. elegans*, *S. pombe*, *T. spiralis*, and *A. thaliana* all strikingly were found to be activated upon N-terminal ubiquitination (Figure 4F). Alphafold3 model predictions of each UCHL5 homolog N-terminally fused to Ub show Ub binding in the back-site (Supplementary Figure 8). Despite representing vastly different kingdoms of life, this exhibition of evolutionary conservation and/or convergence highlights an importance on an allosteric Ub binding site that regulates enzyme activity.

### N-terminal ubiquitination of UCHL5 inhibits debranching activity

As activation towards monoUb substrates of Ub-UCHL5 is achieved by intramolecular association of Ub and the noncanonical Ub binding site, we asked whether this would bias the substrate preference of UCHL5 in the context of larger chain cleavage reaction paths as the back site is indispensable towards debranching activity. UCHL5 is a unique Dub that specifically debranches at Lys48 branch points^59^. This activity was shown to affect degradation rates of substrate decorated with branched Ub architectures and is dependent on F117 and F121 of UCHL5^60^. We hypothesized that the intramolecular Ub association to α5 and α6 of UCHL5 could sterically occlude substrates that would otherwise require that site. As a model system, in vitro Lys6/Lys48 branched Ub trimer cleavage assays show no debranching upon incubation with Ub-UCHL5 compared to the active, unmodified UCHL5 WT control (Figure 5A). This inhibition extends to native-like high molecular weight chains bearing branched ubiquitination, either purely at Lys6/Lys48 or heterogeneously with Lys6-Lys11- and Lys63/Lys48 branched architectures (Figure 5B). Even among N-terminally modified UCHL5 homologs, inhibition of chain debranching is largely conserved relative to their unmodified counterparts, with *C. elegans* and *T. spiralis* exhibiting near-complete inhibition (Supplementary Figure 9C).

**Figure 5.**
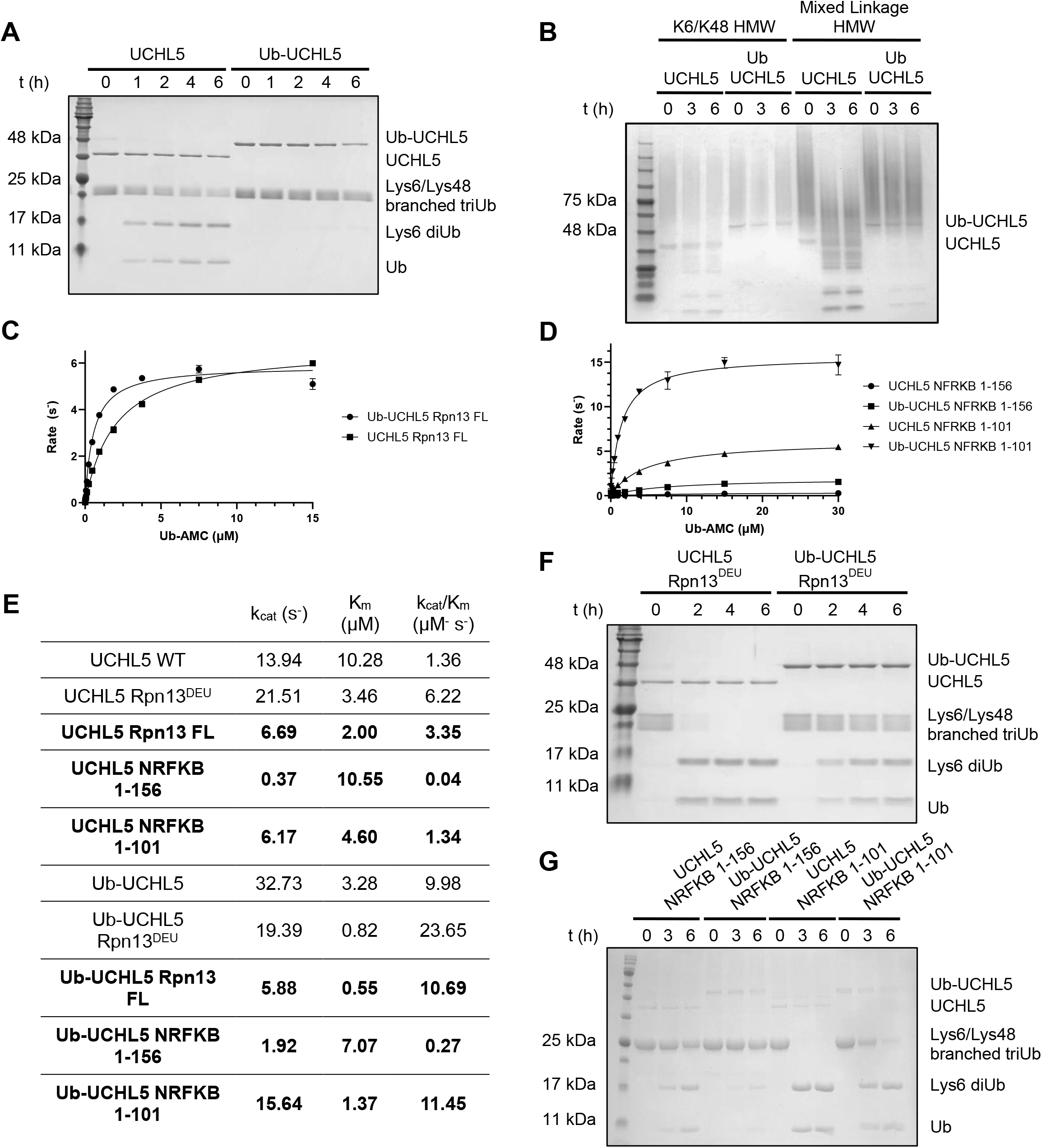
Substrate biasing identified in UCHL5 proteoforms by relief in the inhibition of debranching activity. SDS-PAGE Coomassie-stained gels depict inhibition of debranching by N-terminal ubiquitination of UCHL5 with Lys6/Lys48 branched Ub trimer substrate (A) and high molecular weight Ub chain substrates bearing Lys6/Ly48 and Lys6-, Lys11-, Lys63/Lys48 branch points (B). Michaelis Menten kinetic plots illustrating the effect of N-terminal ubiquitination when in complex with DEUBAD domain containing proteins Rpn13 (C) and NFRKB (D). (E) A table summarizing the Michaelis Menten parameters for the enzymes tested in (C and D). SDS-PAGE Coomassie-stained gels association of Rpn13^DEU^ but not NFRKB^DEU^ relives the inhibition of debranching of the Lys6/Lys48 branched triUb substrate (F and G).

### Rpn13 relieves N-terminally regulated inhibition of UCHL5 debranching activity and suggests debranching being exclusive to the 26S proteasome

Unlike N-terminal ubiquitination which activates C-terminal cleavage of monoUb substrates but inhibits the polyUb chain debranching reaction path of UCHL5, Rpn13 activates UCHL5 in both reaction paths. The activity towards C-terminal cleavage of monoUb substrates is synergistic with these two activation modes. Interestingly different fragments of NFRKB either 1-156 or 1-101 exert either inhibition or activation, respectively, towards Ub-AMC^57^. To further investigate the effect of N-terminal ubiquitination on UCHL5, we reconstituted the modified enzyme in association with Rpn13 and NFRKB fragments to potentially understand their behavior when associated with large protein complexes. Having shown that Rpn13^DEU^ and N-terminal ubiquitination synergistically activate UCHL5, we found that this effect is also observed with Rpn13^FL^ (Figure 5C). As expected, NFRKB^1-156^ inhibited while NRFKB^1-101^ activated Ub-UCHL5 towards Ub-AMC (Figure 5D). N-terminal ubiquitination activated these complexes 3.1, 6.8 and 8.5-fold respectively (Figure 5E). With marginal differences in the K_m_, activation seems to be driven by the *k*_cat_ in the context of the NRFKB 1-156 fragment which is consistent with our previous observations.

We questioned if Rpn13 has bearing on the inhibition of chain debranching path of Ub-UCHL5. Surprisingly, both Rpn13^FL^ and Rpn13^DEU^ are sufficient to relieve the inhibition of debranching activity and reinstate the ability for Ub-UCHL5 to debranch at Lys48 branch points in both the trimer (Figure 5F) and high molecular weight chain architectures. A similar yet less pronounced affect was observed with the NFRKB^1-101^ fragment mediated activation while the NFRKB^1-156^ fragment remained inhibitory (Figure 5G). Taken together, these results suggest that debranching is more pronounced and spatially restricted at the 26S proteasome.

This substrate-dependent behavior raised the possibility that N-terminal ubiquitination alters how UCHL5 engaged Rpn13 and, by extension, the larger 19S regulatory particle. Biolayer interferometry measurements with bait Rpn13^FL^ indicate only a marginal difference in K_D_ between UCHL5 (60 nM) and Ub-UCHL5 (100 nM) to Rpn13^FL^ (Supplementary Figure 10). Alphafold3 predictions of Ub-UCHL5 with Rpn13^DEU^ suggests a shift in the intramolecular association of the N-terminally fused Ub from the back site to a neointerface generated by Rpn13 (Supplementary Figure 11A). This displacement from the back site could be the causative effect in the limiting *k*_cat_ observed in the Ub-AMC assay, much like the abrogation of activation elicited by ncNb displacing the back site intramolecular association. However, alanine mutation along the Rpn13 interface did not affect relief in debranching (Supplementary Figure 11B). Synchrotron-based small-angle X-ray scattering (SAXS) that is particularly sensitive to global structural features of protein assemblies in solutions was used to determine any Rpn13-dependent transitions^82^. SAXS analysis of the Rpn13^DEU^ complex did not support the Alphafold3 neointerface mediate displacement but instead represented a conformation similar to Ub bound in the back side with Rpn13^DEU^ binding to the ULD as previously described^58,72^ (Supplementary Figure 12). Thus, N-terminal ubiquitination does not appear to simply remodel the Rpn13 – UCHL5 interface or alter Rpn13 affinity. Rather, these findings support a model in which the proteasome relieves autoinhibition of Ub-UCHL5 through a context dependent mechanism that enhances debranching only when UCHL5 is properly positioned within the 26S regulatory particle.

## DISCUSSION

In summary, this study identifies substrate biasing in UCHL5 proteoforms. N-terminal ubiquitination of UCHL5, a product of E3-independent noncanonical ubiquitination by UBE2W, confers activation toward cleaving short peptides or other leaving groups from the C-terminus of Ub, consistent with the canonical view of catalytic function of UCH enzymes. This modification, however, inhibits the ability of the enzyme to debranch at Lys48 branch points in solution and potentially rescues substrates from nonspecific debranching until their arrival to the proteasome. Inhibition of debranching is relieved upon association to Rpn13, suggesting debranching activity is spatially restricted at the 26S proteasome. Together, these findings establish a framework in which N-terminal ubiquitination tunes UCHL5 activity differently in solution and in the context of multi-subunit protein complexes. The biochemical, crystallographic, molecular dynamics, and spectroscopic data presented here define the structural logic underlying this proteoform-specific substrate selection.

Analyses of the structure suggest that intramolecular binding of the N-terminal Ub in a unique allosteric Ub binding site controls substrate biasing. This feature seems to be evolutionarily conserved in organisms bearing both UBE2W and UCHL5 homologs. Previously reported regulation of BAP1 by selective DEUBAD ubiquitination also seems to proceed via a similar mechanism of Ub engaging in the back site^79^. In each case of activation, a *k*_cat_ affect has been responsible for enhancement of activity. Despite our efforts, the complete mechanism remains elusive however we suspect that long range motion upon Ub binding to the back site of UCHL5 triggers catalytic preorganization and active site plasticity. Recent spectroscopic studies on the UCH enzyme Yuh1’s N-terminus depicts a dynamic gating mechanism wherein the N-terminus is in an equilibrium between non-gated and gated trajectories threading through and from the crossover loop^83^. Deletions and rigidification of the N-terminal residues abrogate activity in both Yuh1 and UCHL1^84^. Recently, a series of UCHL3 covalent inhibitor bound crystal structures similarly depicts variability in N-terminal orientations^85^. Perhaps, similar gating mechanisms are bypassed with UCHL5 to retain a consistently active form by N-terminal fixation of Ub through backside association.

While the biological circumstances surrounding UBE2W N-terminally ubiquitinating UCHL5 is unknown, its identification by Gly-Gly-Met recognizing antibodies dependent on UBE2W inducible expression, evolutionary conservation, and the innate proximity of the N-terminus to the noncanonical site suggests this modification to be functional rather than incidental. Emerging themes of nonproteogenic ubiquitination^86–89^, Ub fusion degradation (UFD)^90,91^, and defining UCHL5’s role on the INO80 chromatin remodeler complex indicates that many different endogenous substrates could be biased against. Spatiotemporally controlled substrate biasing of Dubs in the UFD pathway suggests that dynamic Dub proteoforms could function as regulatory switches for Ub precursor maturation and chain processing during cellular stress^90^. It is therefore plausible that N-terminal ubiquitination of UCHL5 is a stimulus-dependent phenomenon whose abundance is actively regulated in response to stress rather than a constitutive protein feature. Dub proteoforms, substrate selection, and its effect on proteostasis is an understudied aspect of Dub biology. While only a few Dub proteoforms have been identified, this study, to our knowledge, represents the first to report the molecular basis of proteoform selective substrate biasing.

## METHODS

### Cloning, expression, and purification of recombinant proteins

Expression constructs of His-TEV-Ub-UCHL5 G76V were a generous gift from Genentech. PCR amplified gene fragments of UCHL1 WT, UCHL3 WT, UCHL5 WT, UCHL5-His WT, UCHL5 CD, Ub-UCHL5 G76V, and Rpn13^269–388^ were all cloned into pGEX-6P-1 vector using BamHI or EcoRI as the 5’ restriction enzyme and either NotI or XhoI as the 3’ restriction enzyme followed by ligation. A PCR amplified Ub-His, Ub-UCHL5-His, Ub-UCHL5 CD, Ub-UCHL1, and Ub-UCHL3 gene fragments flanked with cut sites NdeI and XhoI was cloned into an in-house modified pET-28a vector consisting of 5’-CATATG-3’ defining the open reading frame by restriction digest and ligation. All UCHL5 homologs genes, including BAP1, along with their N-terminally ubiquitinated counterparts were gene synthesized and subcloned into the BamHI/XhoI cassette of pGEX-6P-1 (Genscript). Sortase eSrtA (2A-9) expression plasmid was a gift from David Liu (Addgene plasmid #75145). Sortase substrates used in this study were Ub-LALTGGHHHHHH and GG-UCHL5. A Ub-LALTGGHHHHHH gene fragment was cloned into the NdeI/XhoI cassette of pET-21a. The GG-UCHL5 protein was generated by modifying the HRV-3C recognition site^92^ in pGEX-6P-1 harboring UCHL5 CD from LEVLFQGP to LEVLFQGG by site directed mutagenesis. Genes for the cNb and ncNb were cloned into pET-28b supporting a 5’ hexahistidine sequence tag. Expression of NRFKB DEUBAD alone led to a significant portion being observed in inclusion bodies. This complex was expressed together with both inserts being cloned into pETDUET with GST-UCHL5 and Ub-UCHL5 in the first multiple cloning site and NRFKB 1-156 or 1-101 in the second multiple cloning site. The source vector (Addgene plasmid #61939) was a gift from Christopher Hill and cloned accordingly. Ubiquitin^1–75^ was cloned into pTXB1 and expressed as intein-chitin binding domain fusion protein for the generation of Ub glycine vinyl methyl ester. Cloning, expression and purification of proteins for the generation of Ub linkage specific substrates were carried out as previously described^59,60^. Finally, pET-19b harboring the Adrm1 gene bearing an N-terminal decahistidine tag was a gift from Joan Conaway & Ronald Conaway (Addgene plasmid #19423). Plasmid propagation was mediated using chemical competent *E. coli* DH5α cells (Novagen). All subsequent clones and mutations were created using traditional recombinant cloning methods and seamless ligation (Bioneer and New England BioLabs).

All plasmids were sequence verified prior to expression by Sanger and Next Generation Sequencing (Azenta Life Sciences), transformed into *E. coli* BL21 (DE3) cells (Novagen), and plated on Luria Broth agar containing the appropriate amount of antibiotics (100 μg/mL ampicillin or 50 μg/mL kanamycin). Starter cultures were inoculated and grown for 16 hrs. at 37°C. A 1:100 dilution of the starter culture was used to inoculate larger expression cultures. Cells were grown while shaking at 37 °C until an OD_600nm_ = 0.6 – 0.8 was reached where then protein expression was induced by the addition of 0.5 mM IPTG and carried out for 18 – 20 h at 16 – 18 °C. Cells were harvested at 7000 rpm for 10 min using centrifugation. Harvested cell pellets were frozen in liquid nitrogen upon stored at −80 °C until further use.

Cell pellets containing glutathione S-transferase (GST) fusion protein harboring an HRV-3C PreScission protease site were resuspended in 1X phosphate-buffered saline (PBS) pH 7.0 with 400 mM KCl containing 0.5 mg/mL lysozyme and subject to high pressure disruption by French press. The lysed resuspension was clarified by ultracentrifugation at 75,000 x g for 1 h at 4 °C and applied to a gravity chromatography column (Bio-Rad) containing preequilibrated 10 mL of glutathione-agarose resin (ThermoFisher Scientific). Loading proceeded for 1 h prior to washing with 40 CV of 1X PBS pH 7.0 400 mM KCl. The protein was eluted in 4 CV of 1X PBS buffer pH 7.4 containing 15 mM reduced glutathione. Tag cleavage was carried out overnight at 4 °C by the addition of GST-tagged PreScission Protease to the elution as per the manufacturer’s recommendation (GE Healthcare). This mixture is then concentrated and further purified by size exclusion chromatography using an FPLC sporting a HiLoad Superdex75 16/600 column connected to a Akta Pure FPLC (Cytiva). Fractions of interest were pooled and applied to pre-equilibrated glutathione resin to remove excess tag or protease contamination if necessary. Homogeneity was assessed by sodium dodecyl sulfate polyacrylamide gel electrophoresis (SDS-PAGE). Pure protein was concentrated to 10-40 mg/mL for further experimentation.

Cell pellets containing polyhistidine recombinant fusion tags were lysed and clarified as previously described. The clarified lysate was applied to a gravity chromatography column containing 5 mL of Ni-NTA agarose resin (ThermoFisher Scientific). Loading proceeded for 30 min prior to washing with 30 CV of 1X PBS pH 7.0, 400 mM KCl followed by 10 CV of three different imidazole each containing 1X PBS buffers (30 mM, 60 mM, and 90 mM). The protein was eluted in 8 CV of 1X PBS pH 7.4, 300 mM imidazole, concentrated, and further purified by size exclusion chromatography as described above. Homogeneity was assessed by SDS-PAGE, fractions of interest were pooled and concentrated to a final concentration of 10-40 mg/mL for subsequent experiments.

Cell pellets containing expressed Ub^1-75^-intein-CBD linear fusions were resuspended in 50 mM MES pH 6.0 containing 300 mM sodium acetate and lysed by French press. The lysate was clarified as above and applied to a gravity chromatography column containing preequilibrated 50 mL of chitin resin (NEB) for 1 h at 4 °C. After washing with 5 CV lysis buffer, 1.5 CV of lysis buffer containing 50 mM sodium 2-mercaptoethanesulfonate (elution buffer) was added to the column and incubated overnight at 37 °C. The eluate was collected the next day and the column was washed with another 1.5 CV of elution buffer. Protein was concentrated using a 3 kDa Amicon concentrator, flash frozen, and stored at −80 °C until further use. Generation of Ub vinyl methyl ester proceeded in a reaction volume of 10 mL as follows: to 6 mL of 1 M sodium bicarbonate was added 150 mg of N-hydroxysuccinimide, 120 mg of a TFA glycine vinyl methyl ester salt (prepared described previously), and 1.5 mL of the concentrated Ub G75-MESNa thioester. The reaction was diluted to 10 mL using 1 M sodium biocarbonate and incubated at room temperature overnight in the dark. Afterwards, the protein was buffer exchanged and subject to purification by cation ion exchange using FPLC.

### Ubiquitin 7-aminomethyl-4-coumarin and ubiquitin rhodamine 110 assays

Michaelis-Menten kinetics was carried out using Ub 7-aminomethyl-4-coumarin (R&D Systems U550) as the substrate. Assays are performed in black 384 well polystyrene plates (Fisher 12-566-624) with 25 mM Tris pH 7.5, 50 mM NaCl, 0.1% (w/v) BSA, and 5 mM DTT as the assay buffer. Assays were carried out using a final concentration of 1 nM UCHL5, 500 pM UCHL5 Rpn13^269-388^, 500 pM Ub-UCHL5 G76V, 250 pM Ub-UCHL5 G76V Rpn13^269-388^ complex, 500 pM UCHL5 catalytic domain, 125 pM of Ub-UCHL5 G76V catalytic domain, 1 nM UCHL5 INO80G^1-156^ complex, 0.5 nM Ub-UCHL5 G76V INO80G^1-156^ complex, 0.5 nM UCHL5 INO80G^1-101^ complex, 125 pM UbUCHL5 INO80G^1-101^ complex, 1 nM BAP1^1-240^, 0.5 nM Ub-BAP1^1-240^, 2.5 nM UCHL1 and Ub-UCHL1, and 50 pM UCHL3 and Ub-UCHL3. The reaction was initiated by the addition of 2X enzyme to 2X variable Ub 7-aminomethyl-4-coumarin substrate (R&D Systems U-550-050) for a final reaction volume of 20 μL. Substrate stocks were prepared such that the highest final concentration of 30 μM was achieved. A 2-fold serial dilution of the substrate to achieve final concentrations of 15 μM, 7.5 μM, 3.8 μM, 1.9 μM, 938 nM, 469 nM, 234 nM, 117 nM, and 58.6 nM was performed prior to substrate addition. An initial substrate concentration of 20 μM was used for BAP1 CD and Ub-BAP1 CD. The initial substrate concentration used for UCHL1 and UCHL3 along with their modified counterparts was 1 μM. Fluorescence is immediately read with an excitation wavelength λ_EX_ = 345 nm and emission wavelength λ_EM_ = 445 nm on a BioTek Synergy H4 plate reader at intervals of 12 s for 3 h. Reactions were carried out in duplicate and data was processed using Microsoft Excel and GraphPad Prism. Michaelis-Menten parameters were determined by nonlinear regression in GraphPad Prism.

Single concentration substrate activity assays are performed as described above. Assays were carried out using a final concentration of 150 pM enzyme and 300 nM Ub rhodamine 110 substrate (R&D Systems U-555-050). Briefly, in triplicate, to 30 μL of 4/3 X substrate,4X enzyme or enzyme mutant is added. The final volume of this reaction is either 40 μL or 50 μL. Fluorescence is immediately read with an excitation wavelength λ_EX_ = 485 nm and emission wavelength λ_EM_ = 535 nm on a BioTek Synergy H4 plate reader at intervals of 11 s for 1 h. Data was processed and visualized in Microsoft Excel and GraphPad Prism.

Inactivation kinetics and the determination of *k*_inact_/K_I_ was carried out as previously described^93,94^ using the format described above. Briefly, a final concentration of 300 nM Ub rhodamine 100 substrate was added to a two-fold serial dilution of Ub vinyl methyl ester beginning at a concentration of 100 nM and ending at 12 pM. The reaction was initiated by the addition of a final concentration of 150 pM UCHL5, Ub-UCHL5, or UCHL5 Rpn13^DEU^. Fluorescence is immediately read with an excitation wavelength λ_EX_ = 485 nm and emission wavelength λ_EM_ = 535 nm on a BioTek Synergy H4 plate reader at intervals of 11 s. Data was processed using Microsoft Excel and GraphPad Prism. Baseline correction analysis was achieved by correcting for the origin across all samples for further fitting. Each curve was fit to the equation *Y* = *V_o_* × (1 − *e*^−*kobst*^)⁄*k_obs_*. The *k*_inact_/K_I_ was determined by linear regression of derived *k*_obs_ values as a function of inhibitor (Ub vinyl methyl ester) concentration.

### Protein crystallization and data collection

Ub-UCHL5 subcloned into pGEX-6P-1, expressed and purified as described above was used in crystallization trials. Following tag cleavage, protein was further purified on a Superdex S75 column equilibrated in 50 mM HEPES, 200 mM NaCl, pH 7.4, 1 mM DTT and concentrated to 30 mg/ml. This protein crystals were obtained by hanging drop vapor diffusion at 4 °C. Crystals that appeared in 0.20 M lithium citrate containing 20% PEG 3350 diffracted to 2.7 Å at the Stanford Synchrotron Radiation Lightsource on the 12–2 (λ = 0.98) beamline. Data was processed and scaled using HKL3000^95^ in the C222_1_ space group.

### Structure Determination and Refinement

The structure of Ub^G76V^UCHL5 was determined by molecular replacement^96^ using the program PHASER in the Phenix suite^97,98^ by using four copies of UCHL5 (PDB 3IHR) and Ub (PDB 1UBQ) as input models. The segment corresponding to residues in α10-α12 of UCHL5 were deleted from the search model. Model building and refinement were carried out using COOT^99^ and Phenix refine where the current crystallographic R and *R*_free_ values lie at 0.24 and 0.28, respectively. Electron density is well resolved for the majority of the structure with the exception of residues 149-159, 245-256 in UCHL5 for which no electron density was observed. Residues with no clear electron density or poor side-chain density were either left unmodeled or modeled as glycines or alanines. Analysis of the Ramachandran plot indicated that 92.88% of residues fall in the most favored regions, 6.63% of residues fell in the allowed regions, and 0.49% were observed in the disallowed region. A total of 21 water molecules were observed in the asymmetric unit. The final structure of Ub-UCHL5 was validated using MolProbity^100^ and deposited in the Protein Data Bank under the accession code PDB 36HP.

### Molecular Dynamics Simulations

Three systems were studied by all-atom molecular dynamics (MD) simulations: unmodified UCHL5 and two N-terminally ubiquitinated proteoforms, built from either the aligned, active-like (productive) or the misaligned (non-productive) protomers of the N-terminally ubiquitinated UCHL5 crystal structure. Missing loops in the crystal structure were constructed by homology modeling in Molecular Operating Environment (MOE)^101^. Each system was then solvated in explicit TIP3P water, neutralized, and brought to 0.15 M NaCl, and parametrized with the CHARMM36m force field^102^. All-atom simulations were run in NAMD^103,104^ in periodic boundary conditions with particle-mesh Ewald5^105^ for electrostatics, a 12-Å nonbonded cutoff, a 2-fs timestep, and with bonds to hydrogens constrained. After energy minimization and a staged release of harmonic restraints (Table S1), three independent 1-µs replicas of each system were run in the NPT ensemble (310 K, 1 atm), saving the coordinates every 10 ps. Trajectory analyses including the calculation of Cα RMSD and RMSF, back-site and active-site SASA, catalytic-triad and sup-porting active-site distances, N-terminal-Ub/back-site contacts, and residue–residue contact maps were performed in VMD^106^. System construction, the complete simulation parameters, and all analysis definitions are provided in more detail in the Supplementary Methods.

### Isotopic labeling for NMR spectroscopy

The [U-^13^C, ^15^N] ubiquitin-LALTGG-His and Ub-UCHL5-His proteins were overexpressed in *E. coli* BL21(DE3) cells grown at 37 °C in M9 minimal medium supplemented with 1 g/L ^15^NH_4_Cl and 2 g/L [^13^C_6_]-D-glucose as the sole nitrogen and carbon sources, respectively. [U-^2^H, ^13^C,^15^N] GG-UCHL5 CD was expressed in M9 minimal medium prepared in 100% D₂O. All isotopically labeled proteins were purified as described above.

### Sortase-mediated transpeptidation of Ub-UCHL5 CD

Transpeptidation reactions were carried out in reaction buffer containing 50 mM Tris pH 7.5, 150 mM NaCl, and 5 mM CaCl₂. Isotopically labeled or unlabeled GGM-UCHL5 CD and Ub-LALTGG-His were used at final concentrations of 20 µM and 100 µM, respectively, with sortase at a final concentration of 20 µM. The enzyme, substrate, and sortase were each prepared as 3X stock solutions. Reactions were initiated by mixing GGM-UCHL5 CD and Ub-LALTGG-His at a 1:1 (v/v) ratio, followed by the addition of an equal volume of sortase to achieve the final reaction concentrations. The reaction mixture was incubated at 37 °C, and reaction progress was monitored by SDS-PAGE using aliquots collected at defined time points. Upon reaction completion, the mixture was passed through Ni-NTA resin pre-equilibrated with reaction buffer (50 mM Tris, pH 7.5, 150 mM NaCl, 5 mM CaCl_2_) to remove His-tagged components, including unreacted Ub-LALTGG-His and sortase. The flow-through containing the transpeptidation product was concentrated and subjected to SEC on a HiLoad 16/600 Superdex 75 column (Cytiva, USA) equilibrated with 50 mM Tris (pH 7.6), 0.5 mM EDTA, and 0.5 mM TCEP. Peak fractions were collected and purity was confirmed by 12% SDS-PAGE.

### Nuclear Magnetic Resonance Spectroscopy

All NMR spectra were collected using a Bruker NEO 600 MHz with a TCI cryo-probe at 310 K. [U-^13^C, ^15^N] Ub, and UCHL5 CD before and after ligation, and unlabeled Ub ligated with [U-^2^H, ^13^C, ^15^N] UCHL5 CD were prepared in NMR buffer (50 mM Tris (pH 7.6), 0.5 mM EDTA, 0.5 mM TCEP, 0.02 % NaN_3_). For NMR data collection of Ub and its ligated form, [U-^13^C, ^15^N] Ub and [U-^13^C, ^15^N] Ub-UCHL5 CD were concentrated to 500 μM and 300 μM, respectively. The ^15^N-^1^H heteronuclear single quantum coherence (HSQC) spectra and 3D triple resonance spectra, namely HNCA and CBCAcoNH, were collected by non-uniform sampling (NUS) of 35 %, except CBCAcoNH of [U-^13^C, ^15^N] Ub-UCHL5 CD ligation sample was collected at 30 % of NUS. For data collection of UCHL5 CD and its ligated form, [U-^13^C, ^15^N] UCHL5 CD and Ub-[U-^13^C, ^15^N] UCHL5 CD ligation samples were concentrated to 275 μM and 270 μM, respectively. The ^15^N-^1^H HSQC spectra and the 3D backbone triple resonance spectra, namely HNCACB, HNCA, and CBCAcoNH^107^, were collected by NUS of 40 %, 35 %, and 35 %, respectively. For Ub-[U-^2^H, ^13^C, ^15^N] UCHL5 CD ligated sample, the protein concentration was 365 μM. The ^15^N-^1^H transverse relaxation-optimized spectroscopy (TROSY) and the TROSY-based 3D backbone triple resonance spectra, including trHNCACB, trHNCA, and trCBCAcoNH, were collected using NUS with sampling rates of 35 %, 32 %, and 35 % respectively. All NMR spectra were processed by NMRPipe^108^, and the NUS datasets were reconstructed by the SMILE program^109^. The backbone resonance assignments of Ub, Ub after ligation, UCHL5 CD, and UCHL5 CD after ligation were achieved by POKY^110^, and deposited in the Biological Magnetic Resonance Bank (BMRB) under the accession codes 53850, 53851, 53852 and 53853, respectively.

The chemical shift perturbations (CSPs) were calculated by following equation:

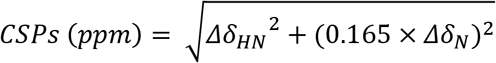

where Δδ_ΗN_ and Δδ_Ν_ are the chemical shift differences between none-ligated and ligated samples for the amide proton and amide nitrogen, respectively. The CSP values were plotted by Prism 11 (GraphPad, USA) and mapped on the crystal structure of Ub-UCHL5 CD using PyMOL v2.4.2 (Schrodinger Scientific, USA).

The samples using the backbone resonance assignments were used in recording of ^15^N backbone relaxation measurements. The experiments were utilized by using a Bruker NEO 600 MHz with a TCI cryo-probe at 310 K. The inversion recovery delays for longitudinal (R_1_) relaxation were set at 100, 200*, 500, 750*, 1250, 1500, and 2000 ms. Carr-Purcell-Meiboom-Gill spin-echo pulses were applied, and the transverse (R_2_) relaxation delays were set at 15.68*, 31.36*, 47.04, 62.72, 78.40*, 94.08, 109.76 ms. The asterisks indicated the technical duplicate in R_1_ and R_2_ experiments. The relaxation rates were calculated by fitting the peak heights in POKY. As conformational exchanges in loop regions may lead to enhanced R_2_ relaxation^111,112^, the R_2_/R_1_ ratios corresponding to the basal levels of individual domains were defined by the residues located within the secondary structural elements observed in the crystal structure of Ub-UCHL5 CD; these R_2_/R_1_ ratios were averaged to obtain an average R_2_/R_1_ ratio of each domain to estimate the overall correlation time. The overall correlation times of individual domains were estimated from the R_2_/R_1_ ratio according to the following equation^113,114^:

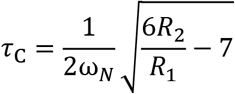

where ω_N_ is the Larmor frequency of ^15^N, which is 60.81 MHz.

### Generation of branched trimer and high molecular weight Ub chain substrates

All branched triUb substrates were made using buffer A (20 mM ATP, 10 mM MgCl2, 40 mM Tris-HCl pH 7.5, 50 mM NaCl, and 1.5 mM DTT). The K6/K48 branched triUb substrate was generated by the addition of 2 mM Ub K6R/K48R, 1 mM Ub D77, 0.5 μM E1, 10 μM UbcH7, 1 μM NleL, and 3 μM AMSH. The K63/K48 and K11/K48 branched triUb substrates were also generated using either 2 mM Ub K48R/K63R, 1 mM Ub D77, 0.5 μM E1, 10 μM cdc34, and 1 μM UbcH13-MmS2 or 2 mM Ub K11R/K48R, 1 mM Ub D77, 0.5 μM E1, 10 μM cdc34, and 20 μM UBE2S-UBD respectively. These reactions were incubated overnight at 37 °C and then quenched by the addition of 50 mM ammonium acetate pH 4.4. All synthesized ubiquitin trimers were further purified from the reaction mixture using size exclusion chromatography (Superdex75) using buffer B (50 mM Tris-HCl pH 7.5, 300 mM NaCl).

High molecular weight Ub chains were generated as previously described^59^. Briefly, both K6-/K48-HMW Ub chains and HMW containing K6-, K11-, K48-, and K63-linkages were assembled in a reaction buffer C (20 mM ATP, 10 mM MgCl2, 40 mM Tris-HCl pH 7.5, 50 mM NaCl, and 0.6 mM DTT) containing 1 mM Ub, 200 nM E1, 5 μM UBE2D3, and 1 μM NleL. For K6/K48 HMW chains 3 μM of AMSH was added 3 hours after the start of the reaction. Both reaction mixtures were incubated at 37°C overnight. Ub chains were purified using size exclusion chromatography (Superdex75) to isolate HMW chains > 75 kDa.

### Lys6/Ly48 triUb and Lys6-, Lys11-, Lys63/Lys48 branched high molecular weight chain debranching assays

Enzymes and substrates were prepared as 2X stocks in the reaction buffer (25 mM HEPES pH 7.5, 2 mM DTT and 50 mM NaCl). The reaction was initiated by mixing the enzyme and substrate at a 1:1 ratio (v/v) to achieve a final concentration of 0.5 μM of the enzyme, 10 μM of Lys6/Ly48 triUb and 0.2 mg/mL of HMW Ub chains. The reaction was incubated at 37°C. Aliquots were collected at the indicated time points and quenched with SDS-PAGE loading buffer.

### Biolayer interferometry

Binding kinetics and affinity measurements were performed using Bio-Layer Interferometry (BLI) on an Octet RED384 system (Sartorius) at 25°C. Ni-NTA biosensors were soaked in assay buffer (PBS, pH 7.4, 0.05% Tween-20, 0.1% BSA) for at least 5 minutes prior to use. His-tagged Rpn13 was loaded onto the biosensor surface at a concentration of 25 µg/mL in assay buffer. The analytes, UCHL5 and Ub-UCHL5, were also prepared in BLI buffer. Association was measured by dipping loaded biosensors into a twofold dilution series of analytes ranging from 1 μM to 7.8 nM. 40 µL of each analyte solution, bait and buffer were added to a 384 tilted-well plate. One Ni-NTA biosensor was used for each K_D_ measurement, dipping the bait protein loaded tip into wells that contained the lowest concentration of the analyte first. The association and dissociation steps were carried out for 120 seconds and 100 seconds, respectively. A reference biosensor loaded with the His-tagged bait protein but dipped into buffer alone was used to correct baseline drift. Biacore Data Analysis Software (version 12.2) was used to collect raw data for the association and dissociation curves. After reference subtraction and baseline alignment in the BAL Octet Data Analysis Software, equilibrium binding responses (R_eq_) were plotted against analyte concentration and fitted to a 1:1 Langmuir binding model to determine the equilibrium dissociation constant (K_D_).

## AKNOWLEDGEMENTS

The authors acknowledge support from the National Institutes of Health NCI and NIGMS (1F31CA275390 to R. P., 5R01 GM126296 to C. D., and R24-GM145965 to E.T.). STDH is supported by Academia Sinica intramural fund, an Academia Sinica Investigator Award (AS-IV-114-L04), and the National Science and Technology Council (NSTC), Taiwan (114-2123-M-001-008). CHL, MKS, and TC are supported by the postdoctoral fellowships from NSTC (114-2811-M-001-106, 114-2811-M-001-116, and 114-2811-B-001-041 respectively). Use of the Stanford Synchrotron Radiation Lightsource, SLAC National Accelerator Laboratory, is supported by the U.S. Department of Energy, Office of Science, Office of Basic Energy Sciences under Contract No. DE-AC02-76SF00515. The SSRL Structural Molecular Biology Program is supported by the DOE Office of Biological and Environmental Research, and by the National Institutes of Health, National Institute of General Medical Sciences (P30GM133894). The contents of this publication are solely the responsibility of the authors and do not necessarily represent the official views of NIGMS or NIH. We thank the Academia Sinica High-field NMR Center (HFNMRC) for the technical support in NMR data collection, funded by the Academia Sinica Core Facility and Innovative Instrument Projects AS-CFII-111-214. We also thank the Protein Core of the Institute of Biological Chemistry, Academia Sinica, for supporting the recombinant protein production, and the staff of the BL13A SAXS beamline at NSRRC.

## AUTHOR CONTRIBUTIONS

This manuscript was written with contributions from all authors. All authors have given approval of the final version of the manuscript. Cloning, protein expression, purification, and biochemical experimentation was performed by R.P., N.P., and D.J. Protein crystallization, data collection, and structure solution was solved by R.P. and N.P. Nuclear magnetic resonance spectroscopy sample preparation, data collection, and evaluation was performed by C.H.L., M.K.S., and S.T.D.H. Molecular dynamics simulations and data evaluation were carried out by P.K., S.A., and E.T. Branched Ub chain substrates were generated by E.S. and E.S. Biolayer interferometry experiments were done and analyzed by N.P. and S.I. SAXS data collection, processing, and interpretation was carried out by Y.S.W., T.C., and S.T.D.H. Material was provided by D.F. R.P., N.P., and C.D. conceived of the ideas and the project described here.

