## Supplemental Information for "Substrate biasing in UCHL5 proteoforms"

Size-exclusion chromatography-coupled small-angle X-ray scattering (SAXS) ........ 5-6
**Supplementary Table 2** ........ **7-8**

### SUPPLEMENTARY RESULTS

#### Size exclusion Chromatography of UCHL5 proteoforms

Previous analytical ultracentrifugation experiments have shown that UCHL5 is a dimer in solution at high concentrations[1]. Analytical size exclusion chromatography of N-terminal modified UCHL5 demonstrates that the modification does not affect dimerization but the association of $Rpn13^{DEU}$ shifts the Ub modified enzyme towards a monomeric state, as previously observed for the UCHL5 alone (Supplementary Figure 3A). The ULD serves an integral role in creating the dimer interface along the substrate binding cleft. Elution profiles suggest that this interface remains intact for dimerization to occur along the ULD of each protomer. Given that the low nanomolar concentrations used in our assays are unlikely to support a substantial dimeric population, it is unlikely that the N-terminal Ub modification activates UCHL5 by altering the monomer-dimer equilibrium.

#### C-terminal cleavage activity of N-terminally modified UCHL5 catalytic mutants

To further probe the activation mechanisms, we measured the rate of Ub-Rho hydrolysis by active site and adjacent area mutants, their N-terminally modified counterparts and $Rpn13^{DEU}$ complexes (Supplementary Figure 2). The catalytic triad mutant D179A remains inactive despite N-terminal ubiquitination, indicating that activation requires an intact catalytic triad. Orthogonally, N-terminal ubiquitination restored activity of the catalytically impaired L181A mutant, located adjacent to the active site to near WT levels, and partially rescued the oxyanion-hole mutant Q82A. Together these results suggest an allosteric activation mechanism relay rather than direct active-site engagement. Enhanced activity of N-terminally ubiquitinated L181A and Q82A in the presence of Rpn13 suggests that activation driven by an N-terminal Ub conjugation is mechanistically distinct from that mediated by Rpn13 association, which is proposed to activate UCHL5 through ULD and crossover loop positioning.

#### SAXS analysis of N-terminally ubiquitinated UCHL5 variants

To further investigate the structural and dynamics features of Ub-UCHL5 CD, we resorted to synchrotron-based small-angle X-ray scattering (SAXS) that is particularly sensitive to global structural features of protein assemblies in solution. Size-exclusion chromatography-coupled multi-angle static light scattering (SEC-MALS) showed a monodispersed and highly symmetric elusion peak for Ub-UCHL5 CD corresponding to a monomeric form (Supplementary Figure 5C). The Kratky plot of Ub-UCHL5 CD exhibited a well-defined peak distribution with minor contributions of flexible structural elements that gave rise to a shallow plateau between q = 0.2 and 0.4 (Supplementary Figure 12D). Comparison of the experimental SAXS data and the theoretical SAXS data generated by FoXS[2] based on the crystal structure of Ub-UCHL5 CD – the missing loops were modeled by SwissModel[3] showed excellent agreements up to q = 0.32, indicating that the relative positioning of ubiquitin and $UCHL5_{CD}$ observed in the crystalline state faithfully recapitulates the solution-state characteristics (Supplementary Figure 12A). In other words, once ligated, Ub and UCHL5 CD form a stably bound assembly that tumbles in solution as a rigid entity, in line with the $^{15}N$ relaxation analysis findings. In the case of Ub-$UCHL5^{FL}$ in complex with $Rpn13^{DEU}$, we used AlphaFold3 to generate five models using default settings. Only one of the five models showed the same positioning of Ub relative to UCHL5 CD as observed in the crystal structure, designated as conformation A; the remaining four models exhibited similar positioning of Ub leaning towards $Rpn13_{DEUBAD}$, designated as conformation B (Supplementary Figure 12C). Using the two distinct conformations as inputs to back-calculated theoretical SAXS profiles showed that conformation A is a much better fit to the experimental SAXS profile compared to conformation B (Supplementary Figure 12B), indicating that the presence of the C-terminal ULD domain in UCHL5 bound to $Rpn13^{DEU}$ may not significantly alter the interaction between the N-terminally fused Ub and UCHL5 CD.

### SUPPLEMENTARY METHODS

#### Details of Simulations

#### System construction

Missing loops in the N-terminally ubiquitinated UCHL5 crystal structure were rebuilt by homology modeling in Molecular Operating Environment (MOE)[4], resulting in fully re-solved *productive* and *non-productive* protomers. All three systems were then assembled with VMD/psfgen[5] employing the CHARMM36m protein topology and force field[6]. In the ubiquitinated systems, residues 1-76 correspond to the N-terminal ubiquitin (Ub), while residues 83-301 correspond to UCHL5, mapping to experimental UCHL5 residues 7-225. In the unmodified system, residues 1-220 correspond to UCHL5, mapping to experimental residues 6-225. Each model was solvated in a rectangular periodic box of explicit TIP3P water with a minimum of 15-Å padding on every side, neutralized, and ionized with 0.15 M NaCl. The resulting solvated systems contained ~$6.5\times10^4$ atoms. Coordinates were centered on the system's center of mass before equilibration.

#### Equilibration and production runs

All MD simulations used NAMD[7,8] with a 2-fs timestep. Bonds to hydrogens were held rigid (rigidBonds all in NAMD) using Shake[9]. Particle mesh Ewald (PME)[10] was used for electrostatics (interpolation order 6, grid spacing 1.0 Å), and a 12- Å real-space cutoff with force-based switching from 10 Å was employed for long-range forces. Langevin thermostat and barostat[11,12] were used to maintain the temperature and pressure at 310 K and 1.01325 atm, respectively. Each replica was first energy-minimized and then equilibrated through a series of re-strained equilibration steps. Positional restraints on the proteins were gradually reduced during these steps and were fully removed before the final production runs. The production simulations were performed in triplicates, each for 1 µs, without restraints.

| **Stage** | **Duration** (steps, *t*) | **Protein/Ligand Restraint Force** (kcal/mol/Å$^2$) |
|---|---|---|
| Minimization | 5k steps | N/A |
| Equilibration 1 | 500k, 1.0 ns | 10.0 |
| Equilibration 2 | 250k, 0.5 ns | 5.0 |
| Equilibration 3 | 250k, 0.5 ns | 2.5 |
| Equilibration 4 | 250k, 0.5 ns | 1.0 |
| Equilibration 5 | 250k, 0.5 ns | 0.5 |
| Equilibration 6 | 250k, 0.5 ns | 0.25 |
| **Production** | 500M, 1 µs | 0.0 |

Supplementary Table 1: Multi-stage equilibration protocol and restraint scheme

#### Analysis

**Trajectory post-processing.** Production trajectories were wrapped and aligned before the analysis. Water and ions were removed, and the processed trajectories were analyzed in VMD. Simulation residue IDs were mapped to the corresponding experimental UCHL5 residue numbering before reporting residue-based data.

In the equations below, $T$ is total time, $\mathbf{r}_i(t)$ is the position of atom $i$ in frame $t$, $\langle\cdot\ \cdot\ \cdot\rangle$ denotes averaging, and $|\cdot|$ is the Euclidean norm.

**$C_\alpha$ RMSD.** $C_\alpha$ RMSD was calculated for UCHL5, selected functional regions, and the N-terminal Ub, after superimposing the coordinates onto the original frame (to remove translational and rotational components). For a selection $S$ containing $N_S$ atoms,

$$\mathrm{RMSD}_S^{C_\alpha}(t) = \sqrt{\frac{1}{N_S}\sum_{i=1}^{N_S}\left|\mathbf{r}_i^{C_\alpha}(t) - \mathbf{r}_i^{C_\alpha}(0)\right|^2} \quad \text{(S1)}$$

**$C_\alpha$ RMSF.** Per-residue $C_\alpha$ fluctuations were calculated from the aligned to the first frame of trajectories as

$$\langle \mathbf{r}_i \rangle = \frac{1}{T}\sum_{t=1}^{T} \mathbf{r}_i\,(t),\ \ \mathrm{RMSF}_i^{C_\alpha} = \sqrt{\frac{1}{T}\sum_{t=1}^{T} \left|\mathbf{r}_i^{C_\alpha}(t) - \langle \mathbf{r}_i^{C_\alpha} \rangle\right|^2} \qquad \text{(S2)}$$

**Solvent-accessible surface area (SASA).** SASA was calculated using a 1.4-Å probe radius. For atom *i*, the exposed surface area was estimated as

$$A_i = 4\pi\left(r_i + r_p\right)^2 \frac{n_i^{\mathrm{exp}}}{M}, \qquad \mathrm{SASA}_G(t) = \sum_{i\in G} A_i\,(t) \qquad \text{(S3)}$$

where *G* is the selected atom group, $r_i$ is the atomic radius, $r_p$ is the probe radius, *M* is the number of surface points, and $n_i^{exp}$ is the number of exposed points. SASA was reported for the back site and the active-site residue groups.

**Active-site distances.** Minimum heavy-atom distances were measured between selected active-site residue pairs. For two atom groups *A* and *B*,

$$d_{AB}^{\mathrm{min}}(t) = \min_{a\in A,\ b\in B} |\mathbf{r}_a(t) - \mathbf{r}_b(t)| \qquad \text{(S4)}$$

The active-like catalytic triad fraction was calculated as

$$f_{\mathrm{active}} = \frac{1}{T}\sum_{t=1}^{T} \mathbb{1}\left[d_{\mathrm{C88-H164}}^{\mathrm{min}}(t) \le 5.0\ \text{Å} \wedge d_{\mathrm{H164-D179}}^{\mathrm{min}}(t) \le 3.5\ \text{Å}\right] \qquad \text{(S5)}$$

where $\mathbb{1}[\cdot]$ is the indicator function.

**Ubiquitin/back-site contacts and COM distances.** Heavy-atom contacts were counted between selected Ub side chains and UCHL5 back-site residues using a 4.5-Å cutoff. The corresponding center-of-mass distance was calculated as

$$\mathbf{R}_X(t) = \frac{\sum_{i\in X} m_i \mathbf{r}_i(t)}{\sum_{i\in X} m_i}, \qquad d_{AB}^{\mathrm{COM}}(t) = |\mathbf{R}_A(t) - \mathbf{R}_B(t)| \qquad \text{(S6)}$$

**Residue-residue contact-map occupancy.** Residue-residue contact maps were calculated using heavy atoms. Two residues were considered in contact when any heavy-atom pair was within the selected cutoff distance. The contact occupancy for replica *r* was

$$O_{ij}^{(r)} = \frac{1}{T}\sum_{t=1}^{T} \Theta\left(r_c - d_{ij}^{\mathrm{min}}(t)\right),\ d_{ij}^{\mathrm{min}}(t) = \min_{a\in i,\ b\in j} |\mathbf{r}_a(t) - \mathbf{r}_b(t)| \qquad \text{(S7)}$$

For each system, the three replicas were averaged, and differences between the productive and non-productive systems were calculated as

$$\bar{O}_{ij}^{\mathrm{state}} = \frac{1}{R}\sum_{r=1}^{R} O_{ij}^{(r)},\ \Delta O_{ij} = \bar{O}_{ij}^{\mathrm{prod}} - \bar{O}_{ij}^{\mathrm{nonprod}},\ R = 3 \qquad \text{(S8)}$$

**Size-exclusion chromatography-coupled with multiangle static light scattering (SEC-MALS) analysis**

The absolute molecular weight of Ub-UCHL5 CD was determined by SEC-MALS as previously described using an Agilent 1260 HPLC system (Agilent, USA) coupled with a multi-angle light scattering detector (DAWN, Wyatt Technology) and a refractive index detector (Optilab, Wyatt Technology). Bovine serum albumin, which was used as a reference standard, Ub-UCHL5$_{CD}$, were prepared at a concentration of 1 mg/mL in 50 mM HEPES, 150 mM (pH 7.4) with 0.02% $NaN_3$ and applied to the SEC column with 10 uL. SEC was carried out using a Bio-SEC 3 liquid chromatography column (Agilent). The refractive index increments (dn/dc) of protein was defined as 0.185 mL/g as inputs for the molecular weight calculations by using the ASTRA 8.0 software (Wyatt Technology).

**Size-exclusion chromatography-coupled small-angle X-ray scattering (SAXS)**

The SAXS data of Ub-UCHL5 CD were collected at the BioSAXS beamline 13A of Taiwan Photon Source (TPS) at the National Synchrotron Radiation Research Center (NSRRC), Taiwan. The BioSAXS beamline 13A integrates the SAXS/UV-Vis-absorption measurement with a high-performance liquid chromatography (HPLC) system. 100 μL of protein samples at a concentration of 12 mg/mL was injected into the HPLC system and separated by BioSEC-3 (300 Å, 4.6×300 mm) (Agilent, USA) at a flow rate of 0.35 mL/min. The flow rate was reduced to 0.15 mL/min before the SAXS data collection to extend the duration of SAXS data collection. The exposure time of each frame was set at two seconds. The SAXS data were collected with a momentum transfer q ranging from 0.006 to 0.908 $Å^{-1}$ for a wavelength of 0.827 Å, 15 keV, using an integrated detecting system comprising an Eiger X 9M SAXS detector (Dectris, Switzerland). An in-house LabVIEW-based software accomplished the data reduction, solvent subtraction, and merging of the SAXS data[13]. The SAXS data were further processed and analyzed using *PRIMUS* in the ATSAS 3.2.1 software package to estimate the $R_g$ and $D_{max}$ value based on the Guinier approximation and the pairwise distance distribution function (PDDF, P(r))[14]. The data were plotted using Prism 11. The SAXS profiles of Ub-$UCHL5_{CD}$ and Ub-UCHL5 (full-length) in complex with $Rpn13_{DEUBAD}$ were deposited in the Small Angle Scattering Biological Data Bank (SASBDB) under the accession codes of SAS7828 and SAS7830, respectively.

| Primer Name | Oligonucleotide Sequence (5'-3') |
|---|---|
| BamHI-UCHL5 Fwd | CATCATCATGGATCCATGACGGGCAATGCC |
| UCHL5-NotI Rev | GTCAGTCACGATGCGGCCGCTTATTTGGTTTCCTGAGC |
| UCHL5-His-TGA-XhoI Rev | ATGCTCGAGTCAATGATGATGATGATGATGTTTGGTTTCCTGAGCTTTCTTTGCGTTCTG |
| HRV3C-GG-UCHL5 Fwd | CGGATCTGGAAGTTCTGTTCCAGGGGGGAATGACGGGCAATGCCGGGGAGTGG |
| HRV3C-GG-UCHL5 Rev | CCACTCCCCGGCATTGCCCGTCATTCCCCCCTGGAACAGAACTTCCAGATCCG |
| UCHL5 Q82A Fwd | CGACTTGACACGATATTTTTTGCTAAGGCCGTAATTAATAATGCTTGTGCTACTCAAGCC |
| UCHL5 Q82A Rev | GGCTTGAGTAGCACAAGCATTATTAATTACGGCCTTAGCAAAAAATATCGTGTCAAGTCG |
| UCHL5 C88A Fwd | GCTAAGCAGGTAATTAATAATGCTGCGGCTACTCAAGCCATAGTGAGTGTGTTAC |
| UCHL5 C88A Rev | GTAACACACTCACTATGGCTTGAGTAGCCGCAGCATTATTAATTACCTGCTTAGC |
| UCHL5 F117A Fwd | GGCGAGACATTATCAGAGTTTAAAGAAGCGTCACAAAGTTTTGATGCAGCTATGAAAGG |
| UCHL5 F117A Rev | CCTTTCATAGCTGCATCAAAACTTTGTGACGCTTCTTTAAACTCTGATAATGTCTCGCC |
| UCHL5 F121A Fwd | GAGTTTAAAGAATTTTCACAAAGTGCGGATGCAGCTATGAAAGGCTTGGCACTGAG |
| UCHL5 F121A Rev | CTCAGTGCCAAGCCTTTCATAGCTGCATCCGCACTTTGTGAAAATTCTTTAAACTC |
| UCHL5 D179A Fwd | CCTGTTAATGGGAGACTGTATGAATTAGCTGGATTAAGAGAAGGACCGATTG |
| UCHL5 D179A Rev | CAATCGGTCCTTCTCTTAATCCAGCTAATTCATACAGTCTCCCATTAACAGG |
| UCHL5 L181A Fwd | GGAGACTGTATGAATTAGATGGAGCGAGAGAAGGACCGATTGATTTAGGTGC |
| UCHL5 L181A Rev | GCACCTAAATCAATCGGTCCTTCTCTCGCTCCATCTAATTCATACAGTCTCC |
| UCHL5 229TAA Fwd | GGCCATTGTGTCTGACAGAAAATAAATATATGAGCAGAAGATAGCAGAGTTAC |
| UCHL5 229TAA Rev | GTAACTCTGCTATCTTCTGCTCATATATTTATTTTCTGTCAGACACAATGGCC |
| NdeI-Ub Fwd | CATCATCATCATATGCAGATCTTCGTGAAAACCCTGACCG |
| UCHL1-His-XhoI Rev | ATGATGATGCTCGAGTTAATGATGATGATGATGATGCGCCGCCTTGCACAGC |
| Ub-UCHL3 Rev | CAGCCAACGTTGACCCTCCATCACGCCACGCAGACGC |
| UCHL3-His-XhoI Rev | ATGATGATGCTCGAGTTAGTGGTGGTGGTGGTGGTGCGCCGCGCTCAGCGCAATC |
| BamHI-Ub Fwd | CATCATCATGGATCCATGCAGATCTTCGTGAAAACCCTGACCGGC |
| UCHL5-XhoI Rev | ATGATGCTCGAGTTATTTGGTTTCTTGCGCTTTCTTCGCGTTCTGCTTTTCTTTCGC |
| Ub I36A Fwd | CCAAGACAAAGAAGGCGCGCCGCCGGATCAGCAACGTC |
| Ub I36A Rev | GACGTTGCTGATCCGGCGGCGCGCCTTCTTTGTCTTGG |
| Ub I44A Fwd | CGGATCAGCAACGTCTGGCGTTTGCGGGCAAGCAGCTG |
| Ub I44A Rev | CAGCTGCTTGCCCGCAAACGCCAGACGTTGCTGATCCG |
| Ub L71A Fwd | CCTGCACCTGGTGGCGCGTCTGCGTGGTG |
| Ub L71A Rev | CACCACGCAGACGCGCCACCAGGTGCAGG |
| Ub-UCHL5 L73A Fwd | CCTGGTGCTGCGTGCGCGTGGTGTTATGAC |
| Ub-UCHL5 L73A Rev | GTCATAACACCACGCGCACGCAGCACCAGG |
| Ub-UCHL5 76A Fwd | GCGTCTGCGTGGTGCCATGACCGGTAACGCGG |
| Ub-UCHL5 76A Rev | CCGCGTTACCGGTCATGGCACCACGCAGACGC |
| Ub-UCHL5 G76 Fwd | GCGTCTGCGTGGTGGCATGACCGGTAACGC |
| Ub-UCHL5 G76 Rev | GCGTTACCGGTCATGCCACCACGCAGACGC |
| Ub-UCHL5 Q82A Fwd | GATACCATCTTCTTTGCGAAAGCCGTGATTAACAACGCGTGCGCG |
| Ub-UCHL5 Q82A Rev | CGCGCACGCGTTGTTAATCACGGCTTTCGCAAAGAAGATGGTATC |
| Ub-UCHL5 C88A Fwd | GTGATTAACAACGCGGCGGCGACCCAAGCGATC |
| Ub-UCHL5 C88A Rev | GATCGCTTGGGTCGCCGCCGCGTTGTTAATCAC |
| Ub-UCHL5 F117A Fwd | CCTGAGCGAGTTCAAAGAAGCGAGCCAAAGCTTTGATGCGG |
| Ub-UCHL5 F117A Rev | CCGCATCAAAGCTTTGGCTCGCTTCTTTGAACTCGCTCAGG |
| Ub-UCHL5 F121A Fwd | CAAAGAATTTAGCCAAAGCGCGGATGCGGCGATGAAGGGTC |
| Ub-UCHL5 F121A Rev | GACCCTTCATCGCCGCATCCGCGCTTTGGCTAAATTCTTTG |
| Ub-UCHL5 D179A Fwd | CTGTATGAGCTGGCCGGTCTGCGTGAAGGC |

| | |
|---|---|
| Ub-UCHL5 D179A Rev | GCCTTCACGCAGACCGGCCAGCTCATACAG |
| Ub-UCHL5 L181A Fwd | GAGCTGGACGGTGCGCGTGAAGGCCC |
| Ub-UCHL5 L181A Rev | GGGCCTTCACGCGCACCGTCCAGCTC |
| UCHL5 226TAA Fwd | CAACCTGATGGCGATTGTTAGCTAACGTAAAATGATCTATGAGCAG |
| UCHL5 226TAA Rev | CTGCTCATAGATCATTTTACGTTAGCTAACAATCGCCATCAGGTTG |
| UCHL5 307TAA Fwd | GACCCTGGCGGAACACTAACAACTGATCCCGCTGG |
| UCHL5 307TAA Rev | CCAGCGGGATCAGTTGTTAGTGTTCCGCCAGGGTC |
| BamHI-Rpn13 287 Fwd | CATCATCATGGATCCGACCTGGCCAGTGTGCTGACGC |
| Rpn13 385-XhoI Rev | ATGATGATGCTCGAGGGCGTTGTTCTGCATGGCTTTGGCAAACGC |
| Rpn13 DEU Q351A Fwd | CTTGGCCTCGGGGGCGCTGGGCCCCCTC |
| Rpn13 DEU Q351A Rev | GAGGGGGCCCAGCGCCCCCGAGGCCAAG |
| Rpn13 DEU L352A Fwd | GGCCTCGGGGCAGGCGGGCCCCCTCATGTG |
| Rpn13 DEU L352A Rev | CACATGAGGGGGCCCGCCTGCCCCGAGGCC |
| Rpn13 DEU L355A Fwd | GGGCAGCTGGGCCCCGCGATGTGCCAGTTCGGTCTG |
| Rpn13 DEU L355A Rev | CAGACCGAACTGGCACATCGCGGGGCCCAGCTGCCC |
| Rpn13 DEU Q358A Fwd | GCCCCCTCATGTGCGCGTTCGGTCTGCCTGCAG |
| Rpn13 DEU Q358A Rev | CTGCAGGCAGACCGAACGCGCACATGAGGGGGC |
| Rpn13 DEU 350TAA Fwd | CAGCCTTGGCCTCGTAACAGCTGGGCCCCCTC |
| Rpn13 DEU 350TAA Rev | GAGGGGGCCCAGCTGTTACGAGGCCAAGGCTG |
| BamHI-Ube2w Fwd | CATCATCATGGATCCATGCTCGAGATGGCGTCAATGC |
| Ube2w-NotI Rev | ATGATGGCGGCCGCTCAACAAGTATCATCATGATACCACCATTTTGTTTTCTTTGG |
| SbfI-GST-Fwd | GGCGCGCCTGCAGGTCATGTCCCCTATACTAGGTTATTGGAAAATTAAGGGCCTTGTGC |
| SbfI-HRV3C-Ub-Fwd | CACCTGCAGGTCCTGGAAGTTCTGTTCCAGGGGCCCATGCAGATCTTCGTGAAAACCCTG |
| UbUCHL5-NotI-Rev | ATGATGGCGGCCGCTTATTTGGTTTCTTGCGCTTTCTTCGC |
| 5GEX | GGGCTGGCAAGCCACGTTTGGTG |
| 3GEX | CCGGGAGCTGCATGTGTCAGAGG |
| T7 promoter | CTCGATCCCGCGAAATTAATACGACTCACTATAGG |
| T7 terminator | CGTTTAGAGGCCCCAAGGGGTTATGCTAG |

**Supplementary Table 2.** Oligonucleotide primer sequences used to clone in the study.

| Recombinantly Expressed Protein | Vector | Insert | Cassette |
|---|---|---|---|
| GST-HRV3C-UCHL5 | pGEX-6P-1 | UCHL5 | BamHI/NotI |
| GST-HRV3C-Ub-UCHL5 G76V | pGEX-6P-1 | Ub-UCHL5 G76V | BamHI/XhoI |
| GST-HRV3C-Ub-UCHL5 1-225 G76V | pGEX-6P-1 | Ub-UCHL5 226TAA | BamHi/XhoI |
| GST-HRV3C-Ub-UCHL5 1-306 G76V | pGEX-6P-1 | Ub-UCHL5 307TAA | BamHI/XhoI |
| GST-HRV3C-Ub-UCHL5 | pGEX-6P-1 | Ub-UCHL5 | BamHI XhoI |
| GST-HRV3C-UCHL1 | pGEX-6P-1 | UCHL1 | BamHI/XhoI |
| GST-HRV3C-UCHL3 | pGEX-6P-1 | UCHL3 | EcoRI/XhoI |
| GST-HRV3C-UCHL5-His | pGEX-6P-1 | UCHL5 | BamHI/XhoI |
| GST-HRV3C-BAP1 1-240 | pGEX-6P-1 | BAP1 | BamHI/XhoI |
| GST-HRV3C-Ub-BAP1 1-240 | pGEX-6P-1 | Ub-BAP1 | BamHI/XhoI |
| GST-HRV3C-MmUCHL5 | pGEX-6P-1 | MmUCHL5 | BamHI/XhoI |
| GST-HRV3C-Ub-MmUCHL5 | pGEX-6P-1 | MmUb-UCHL5 | BamHI/XhoI |
| GST-HRV3C-AtUCH2 | pGEX-6P-1 | AtUCH2 | BamHI/XhoI |
| GST-HRV3C-Ub-AtUCH2 G76V | pGEX-6P-1 | AtUb-UCH2 G76V | BamHI/XhoI |
| GST-HRV3C-CeUbh4 | pGEX-6P-1 | CeUbh4 | BamHI/XhoI |
| GST-HRV3C-Ub-CeUbh4 G76V | pGEX-6P-1 | CeUb-Ubh4 G76V | BamHI/XhoI |
| GST-HRV3C-XlUCHL5 | pGEX-6P-1 | XlUCHL5 | BamHI/XhoI |
| GST-HRV3C-Ub-XlUCHL5 G76V | pGEX-6P-1 | XlUb-UCHL5 G76V | BamHI/XhoI |
| GST-HRV3C-TsUCHL5 | pGEX-6P-1 | TsUCHL5 | BamHI/XhoI |
| GST-HRV3C-Ub-TsUCHL5 G76V | pGEX-6P-1 | TsUb-UCHL5 G76V | BamHI/XhoI |
| GST-HRV3C-PfUCH54CD | pGEX-6P-1 | PfUCH54 CD | BamHI/XhoI |
| GST-HRV3C-Ub-PfUCH54 CD G76V | pGEX-6P-1 | PfUb-UCH54 CD G76V | BamHI/XhoI |
| GST-HRV3C-Rpn13 269-388 | pGEX-6P-1 | Adrm1 269-388 | BamHI/XhoI |
| GST-HRV3C-Rpn13 269-349 | pGEX-6P-1 | Adrm1 269-349 | BamHI/XhoI |
| GST-HRV3C-Rpn13 287-385 | pGEX-6P-1 | Adrm1 287-385 | BamHI/XhoI |
| His-Adrm1 FL | pET-19b | Adrm1 FL | XhoI/KpnI |
| UbiquitinG75-Intein-CBD | pTXB1 | Ubiquitin | NdeI/SpeI |
| Ubiquitin-His | pET-28a | Ubiquitin | NdeI/XhoI |
| Ubiquitin-UCHL5-His G76V | pET-28a | Ub-UCHL5 G76V | NdeI/XhoI |
| Ubiquitin-UCHL5 1-228-His G76V | pET-28a | Ub-UCHL5 1-228 G76V | NdeI/XhoI |
| Ubiquitin-UCHL1-His G76V | pET-28a | Ub-UCHL1 | NdeI/XhoI |
| Ubiquitin-UCHL3-His G76V | pET-28a | Ub-UCHL3 | NdeI/XhoI |
| His-cNb | pET-28b | Nb-cS1 | NcoI/XhoI |
| His-ncNb | pET-28b | Nb-ncS1 | NcoI/XhoI |
| His-TEV-Ub-UCHL5 G76V | Genentech | Ub-UCHL5 G76V | NsiI/EcoRI |
| His-TEV-Ub-UCHL1 G76V | Genentech | Ub-UCHL1 G76V | NsiI/EcoRI |
| UCHL1-His | Genentech | UCHL1 | N/A |
| eSrtA (2A-9) | pET-29b | eSrtA | NdeI/BamHI |
| Ubiquitin LALTGG-His | pET-21a | UbLALTGG | NdeI/XhoI |
| Ubiquitin LLALTGG-His | pET-21a | UbLLALTGG | NdeI/XhoI |
| Ub-UCHL5-His | pET-29a | Ub-UCHL5 | NdeI/XhoI |
| His-GST-Ube2w | pET-28a | Ube2W | BamHI/NotI |
| His-GST-HRV3C-UCHL5 NFRKB 1-156 | pETDUET | UCHL5 and NFRKB | SbfI/NotI |
| His-HRV3C-Ub-UCHL5 G76V NFRKB 1-156 | pETDUET | Ub-UCHL5 and NFRKB | SbfI/NotI |

**Supplementary Table 3.** List of plasmids and constructs used in this study.

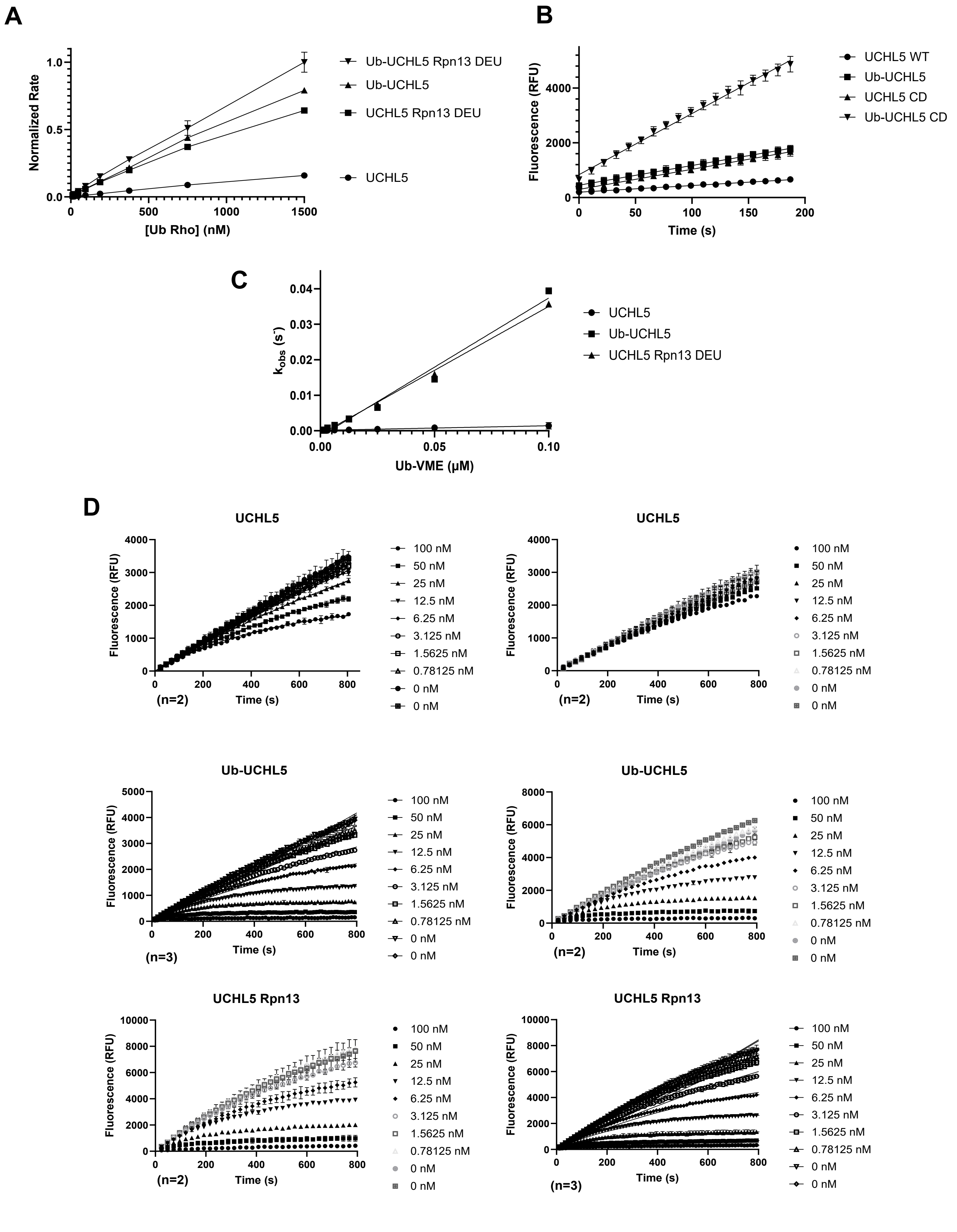
A
Normalized Rate
[Ub Rho] (nM)
Ub-UCHL5 Rpn13 DEU
Ub-UCHL5
UCHL5 Rpn13 DEU
UCHL5
B
Fluorescence (RFU)
Time (s)
UCHL5 WT
Ub-UCHL5
UCHL5 CD
Ub-UCHL5 CD
C
kobs (s-)
Ub-VME (µM)
UCHL5
Ub-UCHL5
UCHL5 Rpn13 DEU
D
UCHL5
Ub-UCHL5
UCHL5 Rpn13
Fluorescence (RFU)
Time (s)
(n=2)
(n=3)
100 nM
50 nM
25 nM
12.5 nM
6.25 nM
3.125 nM
1.5625 nM
0.78125 nM
0 nM
0 nM

**Supplementary Figure 1.** A. C-terminal cleavage is activated by N-terminal ubiquitination of UCHL5 and is synergistic with the activation imparted by Rpn13 DEUBAD using the Ub Rho substrate. B. Activation imparted by N-terminal ubiquitination is independent of the C-terminal ULD domain. Ub Rho cleavage progress curves (top) and the ratio of the initial rates in determining fold activation (bottom). C. N-terminal ubiquitination confers similar cysteine reactivity in UCHL5 than the activation mode elicited by Rpn13 DEU as determined by $k_{inact}/K_i$ measurements using the Ub Rho substrate. D. Raw kinetic progress curves for UCHL5, Ub-UCHL5, and UCHL5 Rpn13. They were fit using a nonlinear regression model to determine $k_{obs}$

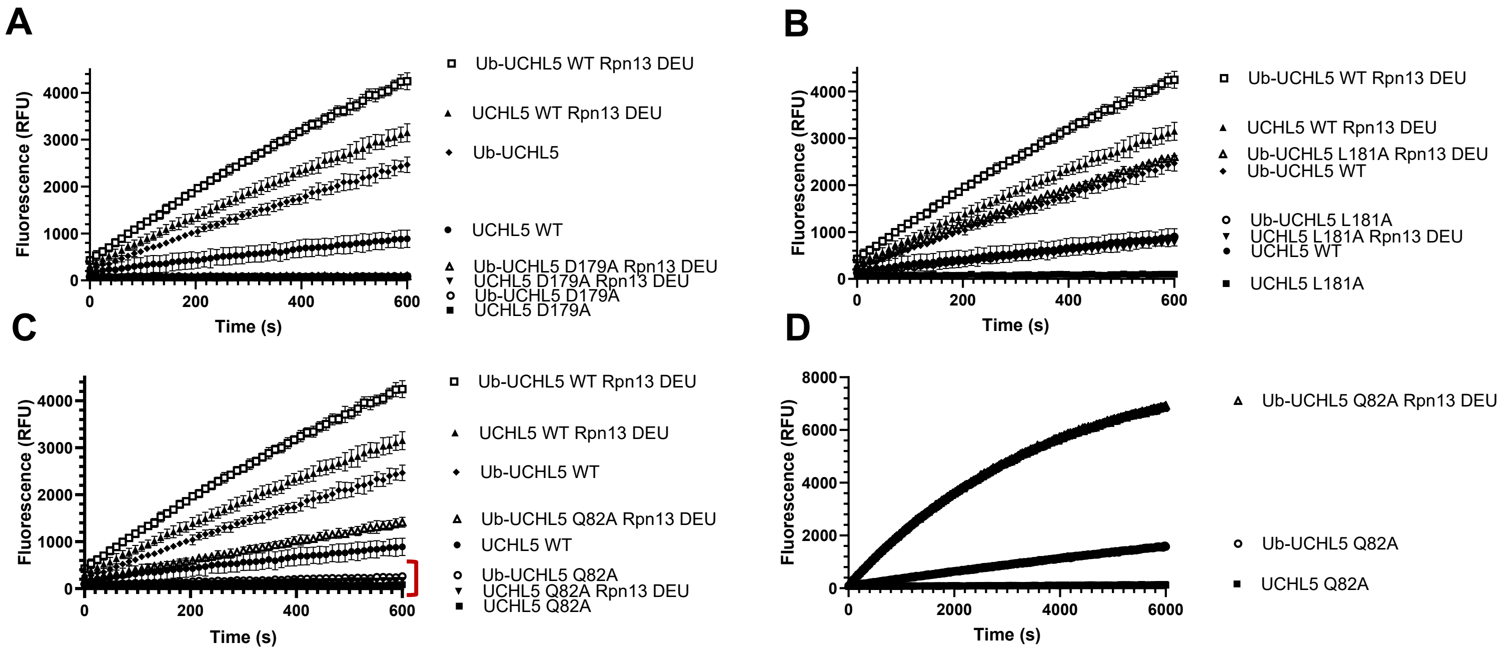

**Supplementary Figure 2.** A. C-terminal cleavage activity of the Ub-Rho substrate by N-terminally modified UCHL5 is perturbed by mutation of the catalytic triad B. N-terminal ubiquitination activates the catalytically inactive L181A mutant suggesting an allosteric mechanism C. D. The activity of oxyanion hole mutation is partially rescued by N-terminal ubiquitination. Activation by Rpn13 association is mechanistically distinct from activation by N-terminal ubiquitination as evidenced by the enhanced cleavage rates observed in the presence of Rpn13.

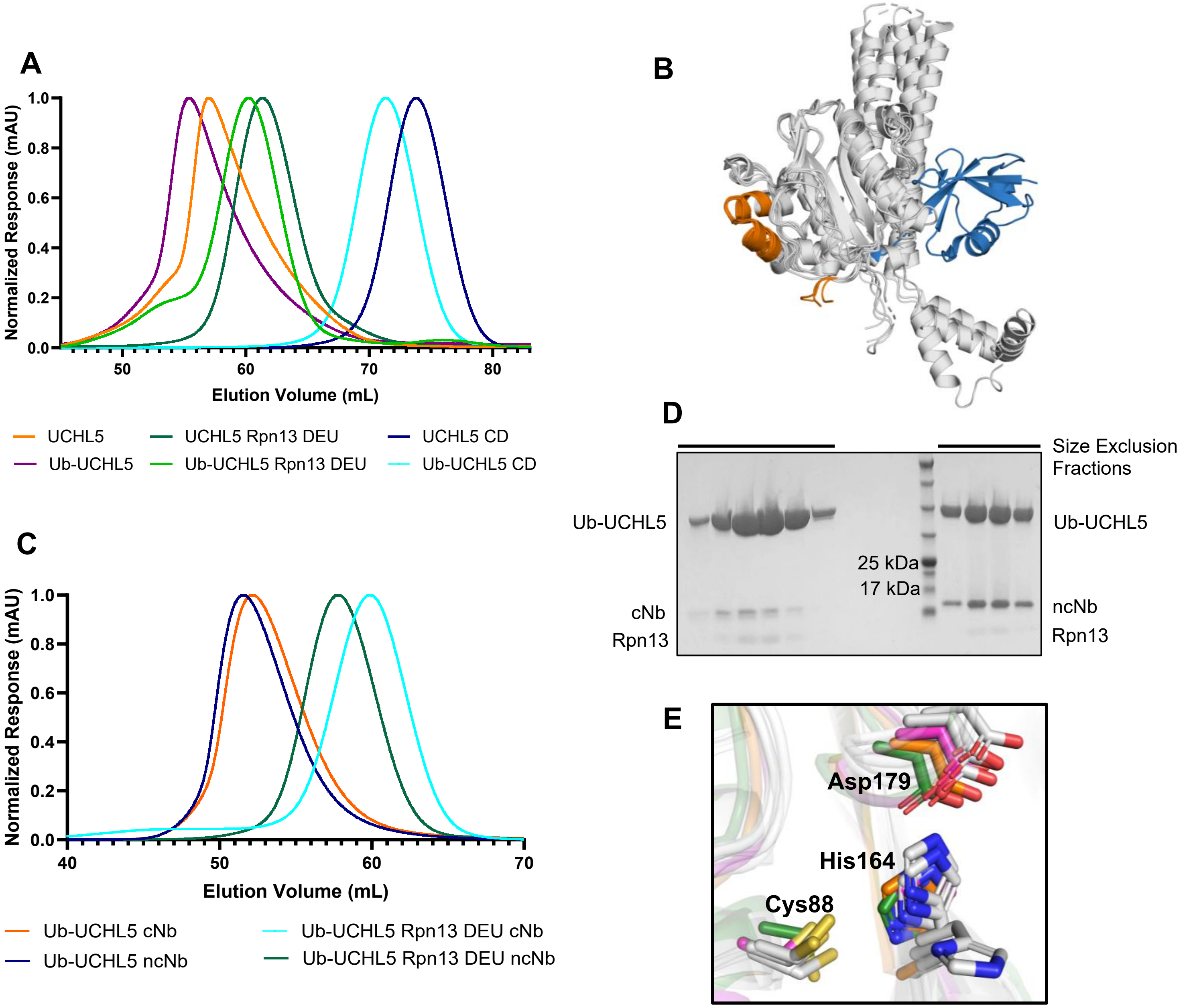

**Supplementary Figure 3.** A. Size exclusion chromatograms of UCHL5 proteoforms depict that N-terminal ubiquitination does not affect dimerization, whereas association with Rpn13 DEUBAD shifts both WT and N-terminally ubiquitinated UCHL5 toward a monomeric state. B. Overlay of UCHL5 crystal structures in the Protein Data Bank (PDB 3RII, 3IHR, 4UEM, 4UEL, 4UF5, 9E7K) reveals that the N-terminus extends towards the "back-site" presented by α5 and α6 helices of UCHL5. (orange) C. Both cNb and ncNb coelute with Ub-UCHL5 and Ub-UCHL5 Rpn13 complex. D. SDS-PAGE Coomassie-stained electrophoresis gel of size exclusion chromatography fractions demonstrating Ub-UCHL5 Rpn13 DEU cNb coelution (left) and ncNb coelution (right). E. Overlay of the catalytic triad of the solved crystal structure of N-terminally modified UCHL5 (gray) with the apo UCHL5 structures (magenta, PDB code: 3IHR; orange, PDB code: 3RII) and the substrate-bound UCHL5 structure (green, PDB code: 4UEL.

| Data Collection | |
|---|---|
| PDB Accession Code | 36HP |
| X-ray Source | SSRL BL 12-2 |
| Wavelength (Å) | 0.98 |
| Resolution range (Å) | 42.17 – 2.735 (2.77-2.73) |
| Space group | $C222_1$ |
| Cell dimensions | |
| a, b, c (Å) | 138.92, 203.98, 149.97 |
| α, β, γ (˚) | 90, 90, 90 |
| Total reflections | 105881(10609) |
| Unique reflections | 54929 (522) |
| Redundancy | 6.2 (6.5) |
| Completeness (%) | 97.31 (28.40) |
| $R_{meas}$ | 0.112 (2.109) |
| $R_{pim}$ | 0.032 (0.592) |
| I/σ(I) | 25.37 (1.52) |
| CC1/2 | 0.995 (0.482) |
| Wilson B ($Å^2$) | 84.49 |
| **Refinement** | |
| Copies/A.S.U. | 4 |
| Resolution (Å) | 2.73 |
| Rwork / Rfree | 0.24 / 0.28 |
| No. nonhydrogen atoms | 11155 |
| Protein | 11155 |
| Ligand | 13 |
| Water | 21 |
| B factor ($Å^2$) | 99.71 |
| Protein | 99.73 |
| Ligand | 123.51 |
| Water | 76.53 |
| R.m.s.d | |
| Bond lengths (Å) | 0.007 |
| Bond angles (˚) | 1.02 |
| Ramachandran (favored/allowed/outliers) | 92.88/6.63/0.49 |
| Clash score | 10.65 |

**Supplementary Table 4.** Data collection, processing, and refinement statistics for the X-ray crystal structure of Ub bound in the noncanonical, allosteric site of UCHL5.

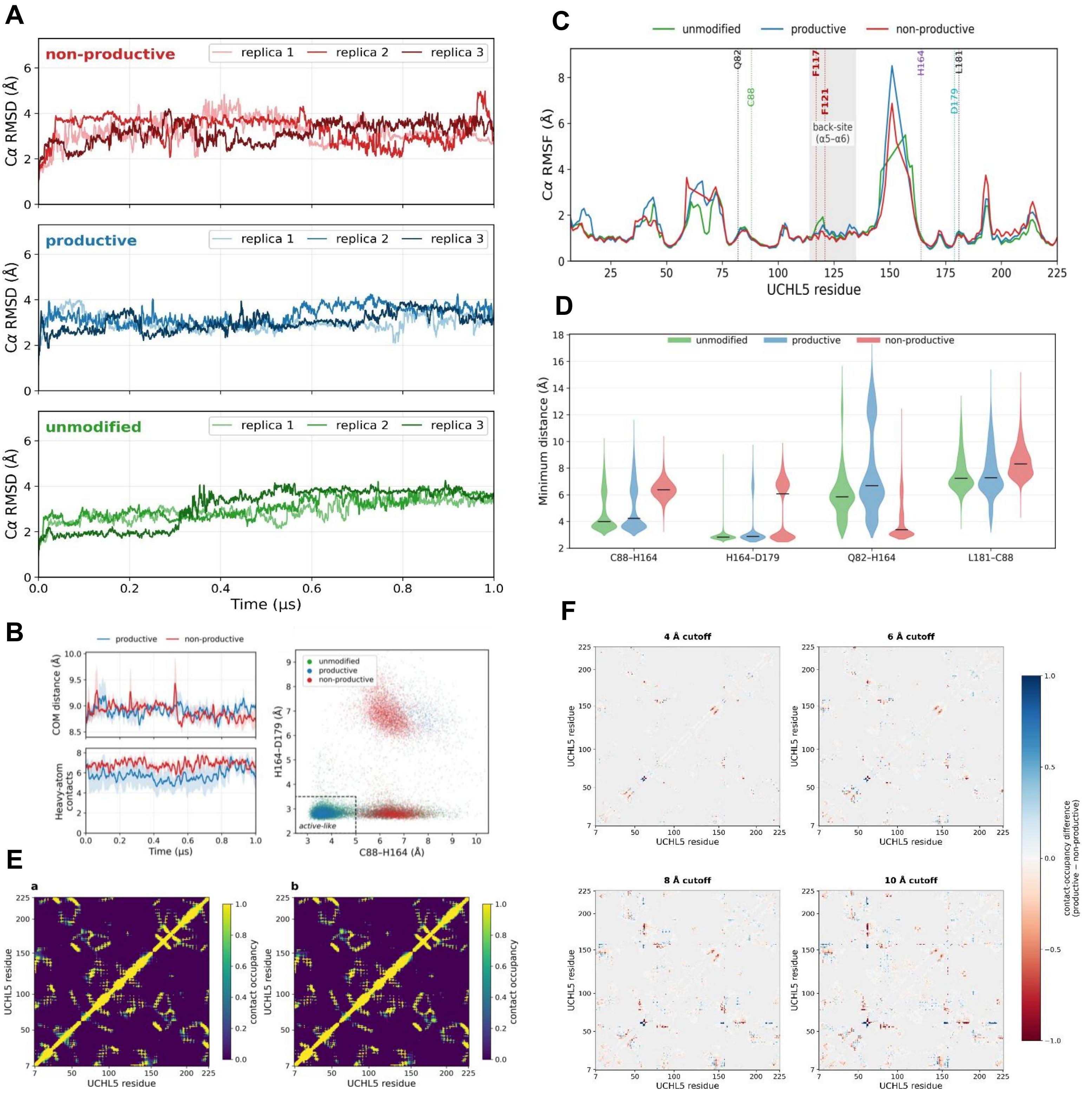

**Supplementary Figure 4.** MD simulations of N-terminally ubiquitinated UCHL5 proteoforms A. Stability of UCHL5 in all simulations. UCHL5 $C_\alpha$ RMSD versus time over the first 1 $\mu$s, plotted for the three replicas of each system. The RMSD rises during early relaxation and then remains bounded around 3–4 Å, with modest replica-dependent drift or transient excursions, showing that the protein remains structurally stable in all three systems. B. Left: N-terminal Ub I36/I44–UCHL5 F117/F121 engagement over the first 1 $\mu$s: mean ± SEM of the center-of-mass distance (top) and heavy-atom contacts (bottom) over three replicas, for both the productive (blue) and non-productive (red) proteoforms; the interface is maintained throughout. Right: Catalytic-triad state based on per-frame closest heavy-atom distances between C88–H164 and H164–D179 from the three replicas per system; the dashed box marks the active-like region (C88–H164 ≤ 5.0 Å and H164–D179 ≤ 3.5 Å), populated mainly by the unmodified and productive proteoforms, and only rarely by the non-productive proteoform (~65%, ~57%, and ~0.7% of frames, respectively). C. Per-residue $C_\alpha$ RMSF. Mean $C_\alpha$ RMSF for the unmodified, productive, and non-productive systems. The back-site $\alpha5$–$\alpha6$ segment (residues 114–135) is shaded, with F117 and F121 marked with red dotted lines. The catalytic triad residues, C88 (green), H164 (purple), and D179 (teal), are marked, and the active-site-peripheral residues, Q82 and L181, are marked in black. The shaded back-site segment, including F117 and F121, shows low $C_\alpha$ RMSF in all three systems, whereas Q82 and L181 are highlighted because they are experimentally implicated residues near the active site and were evaluated together with the active-site distance analyses. D. Active-site closest heavy-atom distance distributions. Distributions of the per-frame shortest heavy-atom distance between each indicated residue pair: the catalytic-triad pairs, C88–H164 and H164–D179, and the supporting pairs, Q82–H164 and L181–C88. Distances were calculated over the three simulation replicas of each system. The non-productive ensemble is shifted toward larger C88–H164 and L181–C88 distances, whereas Q82–H164 is most extended in the productive ensemble, indicating distinct active-site-peripheral rearrangements in the two ubiquitinated proteoforms. These distance distributions underline the catalytic-triad state space shown in B (right) E. Residue–residue contact maps (8 Å cutoff). Residue–residue heavy-atom contact occupancy for (left) productive and (right) non-productive systems, averaged over three replicas across the 198 UCHL5 residues common to both ubiquitinated systems. The two maps are nearly identical (both proteoforms keep the same global contact pattern), so their productive-minus-non-productive difference (F) is localized to a set of residue pairs, mostly in the active-site neighborhood. F. Difference in contacts calculated using different cutoffs. Mean contact differences at the 4, 6, 8 and 10 Å heavy-atom cutoffs, calculated as Productive − non-productive and colored as indicated in the color bar (zero-difference pairs in gray; blue, productive-enriched; red, non-productive-enriched). A value of +1 indicates contacts present only in the productive state, while −1 indicates contacts present only in the non-productive state. Larger cutoffs include more contacts and hence more nonzero pairs, but the localized difference pattern persists across the plotted cutoffs, confirming that the comparison is not an artifact of the contact definition.

**A**

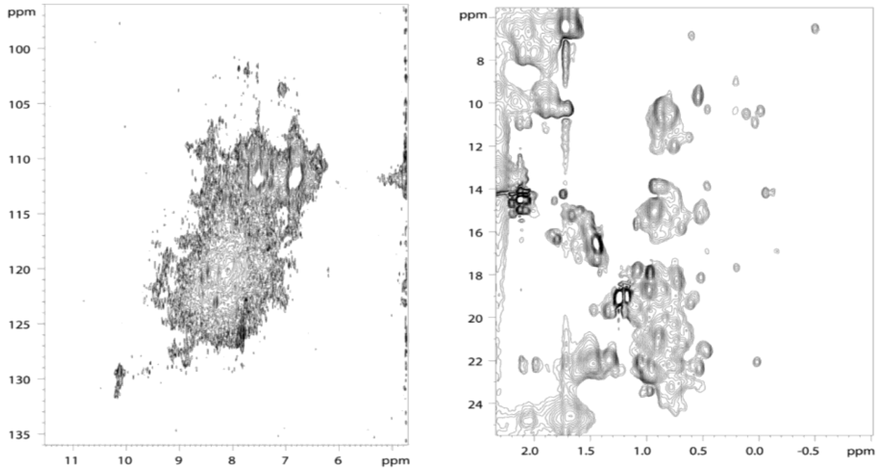

**B**

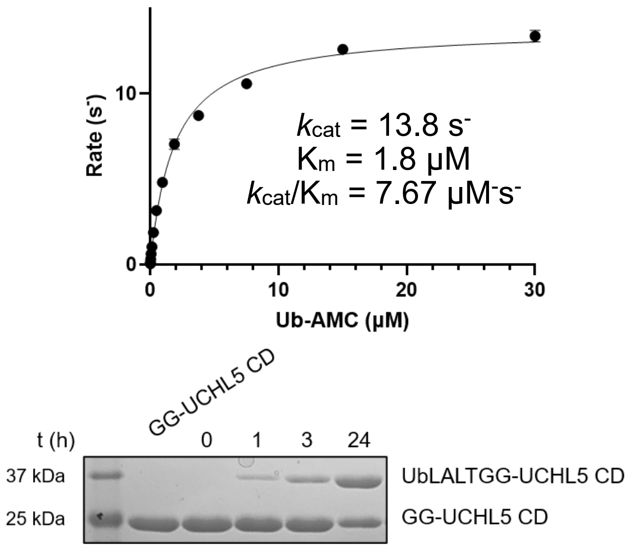

**C**

| Protein | Sortase ligation | N[a] | $R_2/R_1$ | $\tau_C$ (ns) |
|---|---|---|---|---|
| Ubiquitin | Before | 34 | 2.52 ± 0.27 | 3.71 ± 0.36 |
| Ubiquitin | After | 35 | 17.93 ± 4.14 | 13.02 ± 1.63 |
| $UCHL5_{CD}$ | Before | 62 | 10.90 ± 2.64 | 9.93 ± 1.20 |
| $UCHL5_{CD}$ | After | 64 | 18.59 ± 4.47 | 13.28 ± 1.66 |

**D**

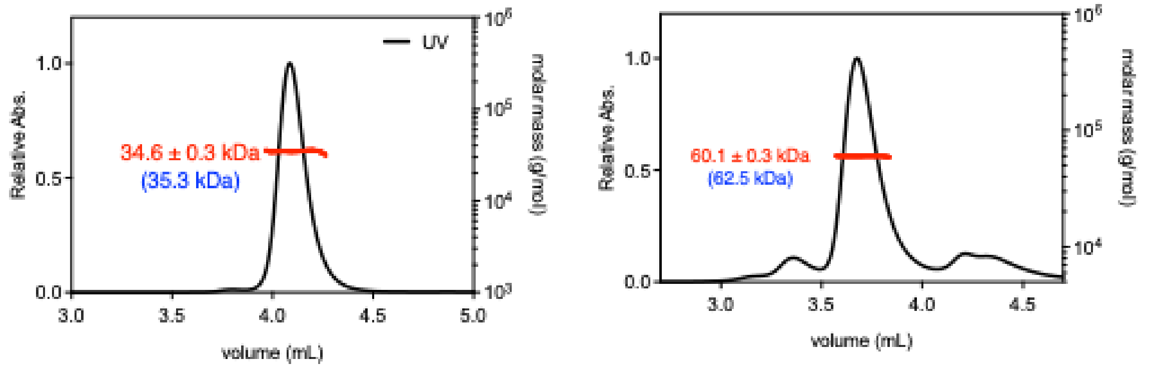

**Supplementary Figure 5.** A. 2D $^{15}$N-$^{1}$H and $^{13}$C-$^{1}$H HSQC spectra of [U-$^{13}$C/$^{15}$N] full-length UCHL5 fused with ubiquitin at the N-terminus. The $^{15}$N-$^{1}$H HSQC spectrum exhibited severely broadened resonances (left). The methyl region of the $^{13}$C-$^{1}$H HSQC spectrum showed some well-dispersed resonances, but the spectral overlaps remain challenging (right). B. Michaelis Menten kinetics of sortase-mediated ligation product of Ub-LALTGG-UCHL5 CD (top) and an SDS-PAGE Coomassie-stained gel of an unoptimized ligation reaction (bottom). C. Table summarizing $^{15}$N spin relaxation $R_2/R_1$ ratios and the corresponding rotational correlation time of ubiquitin and UCHL5 CD (a. Number of residues within the secondary structures included in the $R_2/R_1$ analysis). D. SEC-MALS analysis of Ub-UCHL5 CD (left) and Ub-UCHL5 (full-length in complex with Rpn13$^{DEU}$, right). The SEC UV chromatograms are shown in solid black lines and the MALS-derived molecular mass distributios are shown in red line with the estimated molecualr mass indicated in red labels and the theoretical values indicated in blue parentheses.

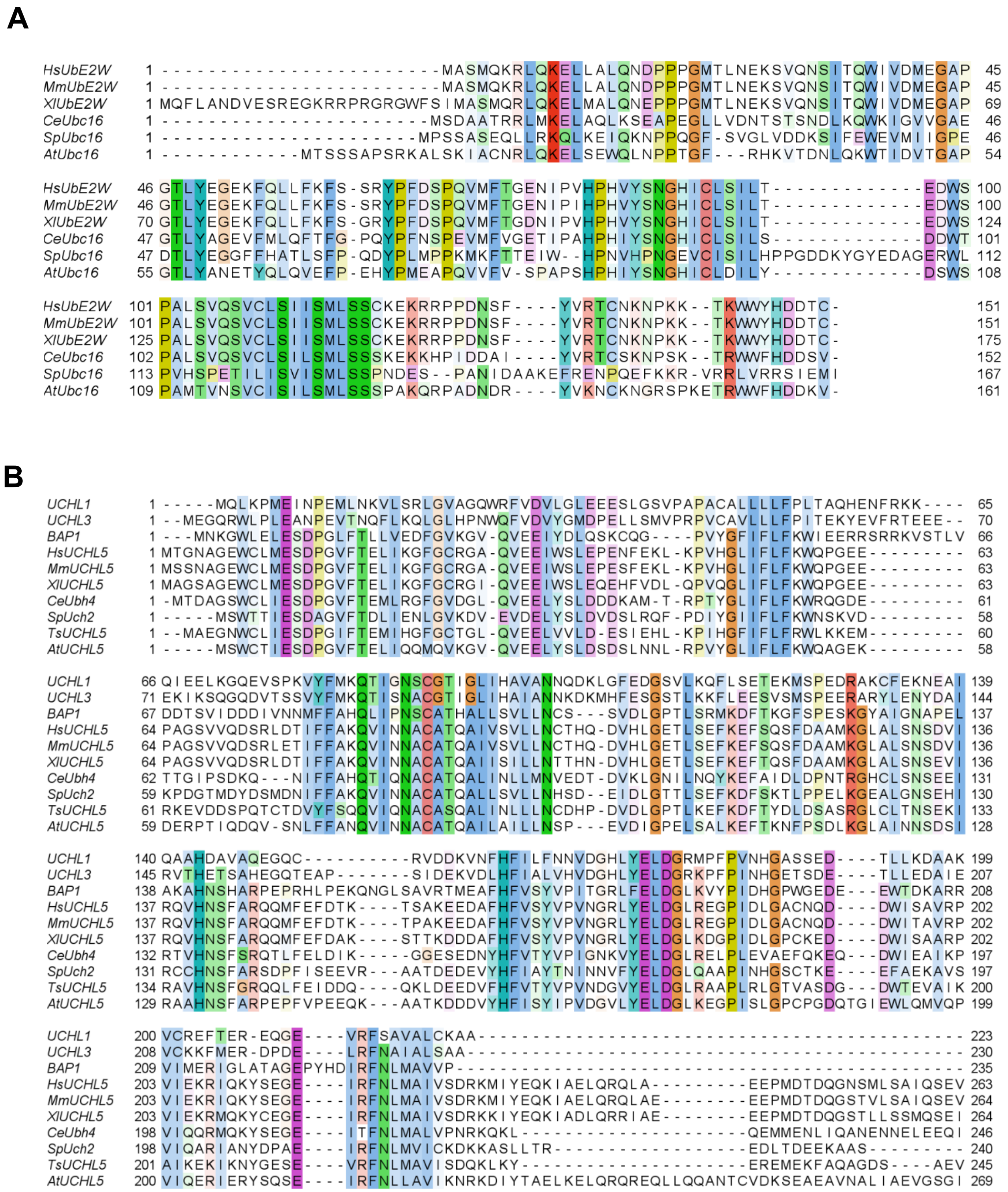

**Supplementary Figure 6.** A. Multiple sequence alignment (MSA) of UBE2W homologs across evolutionary distant species extending to yeast, highlighting sequence conservation and identifying UBC16 as the closest ortholog. B. MSA of UCHL5 homologs across diverse evolutionary lineages highlighting conserved residues in the UCH domain

**A**

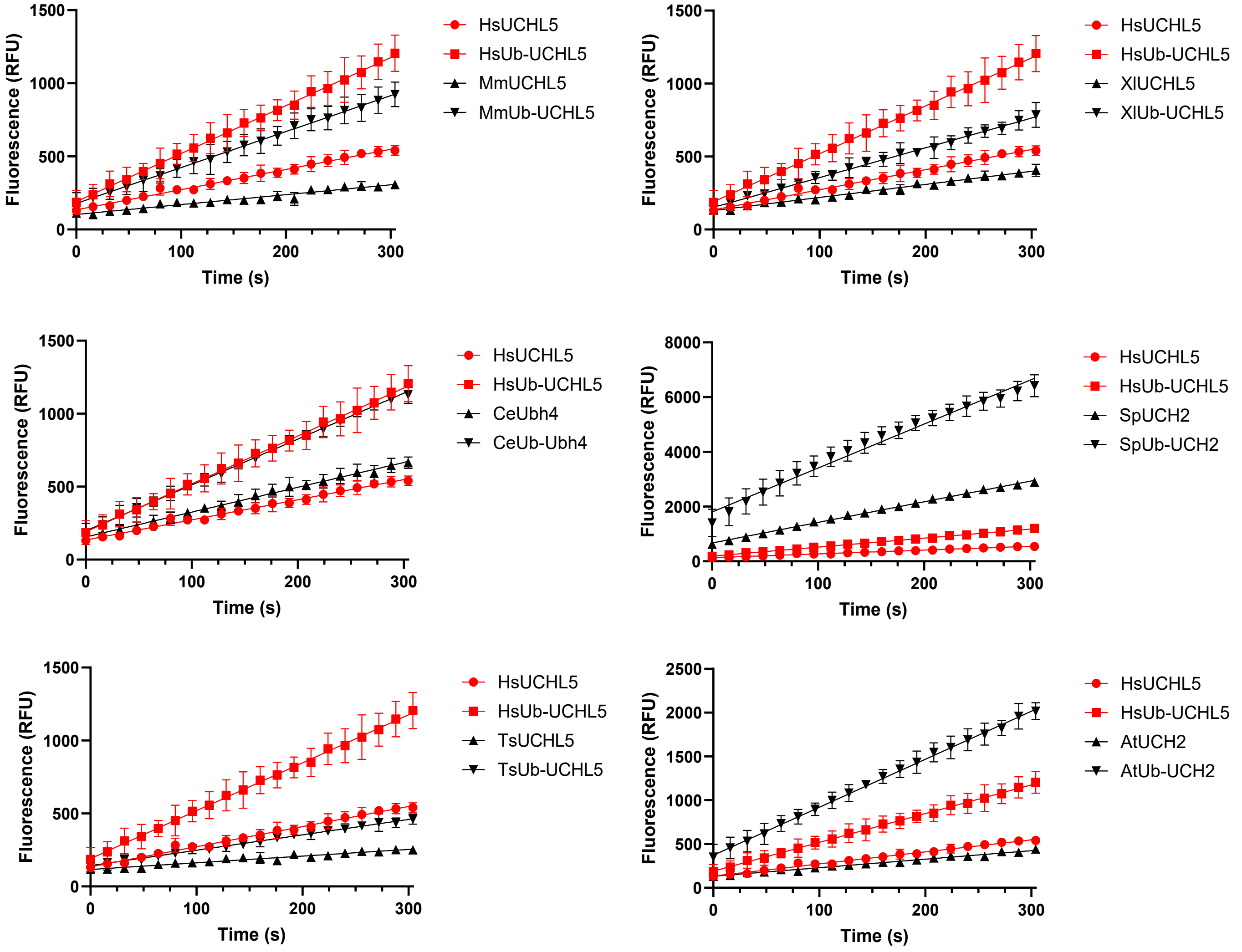

**B**

| | UCHL1 | UCHL3 | BAP1 | Hs | Mm | Xl | Ce | Sp | Ts | At |
|---|---|---|---|---|---|---|---|---|---|---|
| Sequence Identity | 22% | 28% | 44% | - | 97% | 85% | 46% | 40% | 46% | 48% |
| Fold Activation | - | - | 2.66 | 2.38 | 3.64 | 2.28 | 1.85 | 2.13 | 2.30 | 5.63 |

**Supplementary Figure 7.** A. Activation imparted by N-terminal modification is evolutionarily conserved, as reflected by increased fold activation in Ub-Rho assays of UCHL5 homologs and their N-terminally modified counterparts. B. A table summarizing sequence identity relative to human UCHL5 (Hs) and corresponding fold activation upon N-terminal modification.

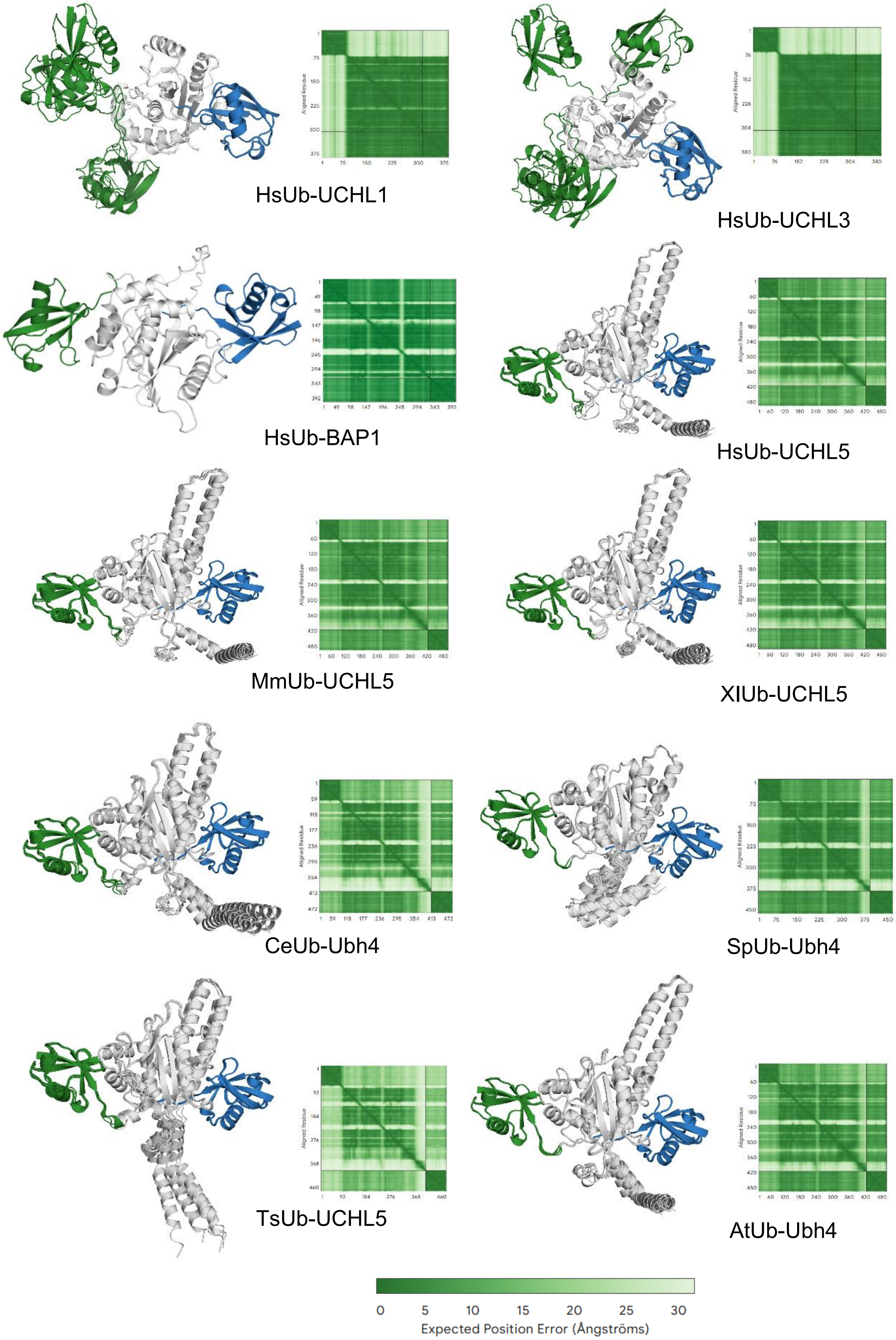

**Supplementary Figure 8.** AlphaFold3 models and Predicted Aligned Error (PAE) plots of UCHL5 homologs across diverse evolutionary lineages reveal conserved N-terminal Ub binding at the back-site (noncanonical Ub:green/canonical Ub:blue).

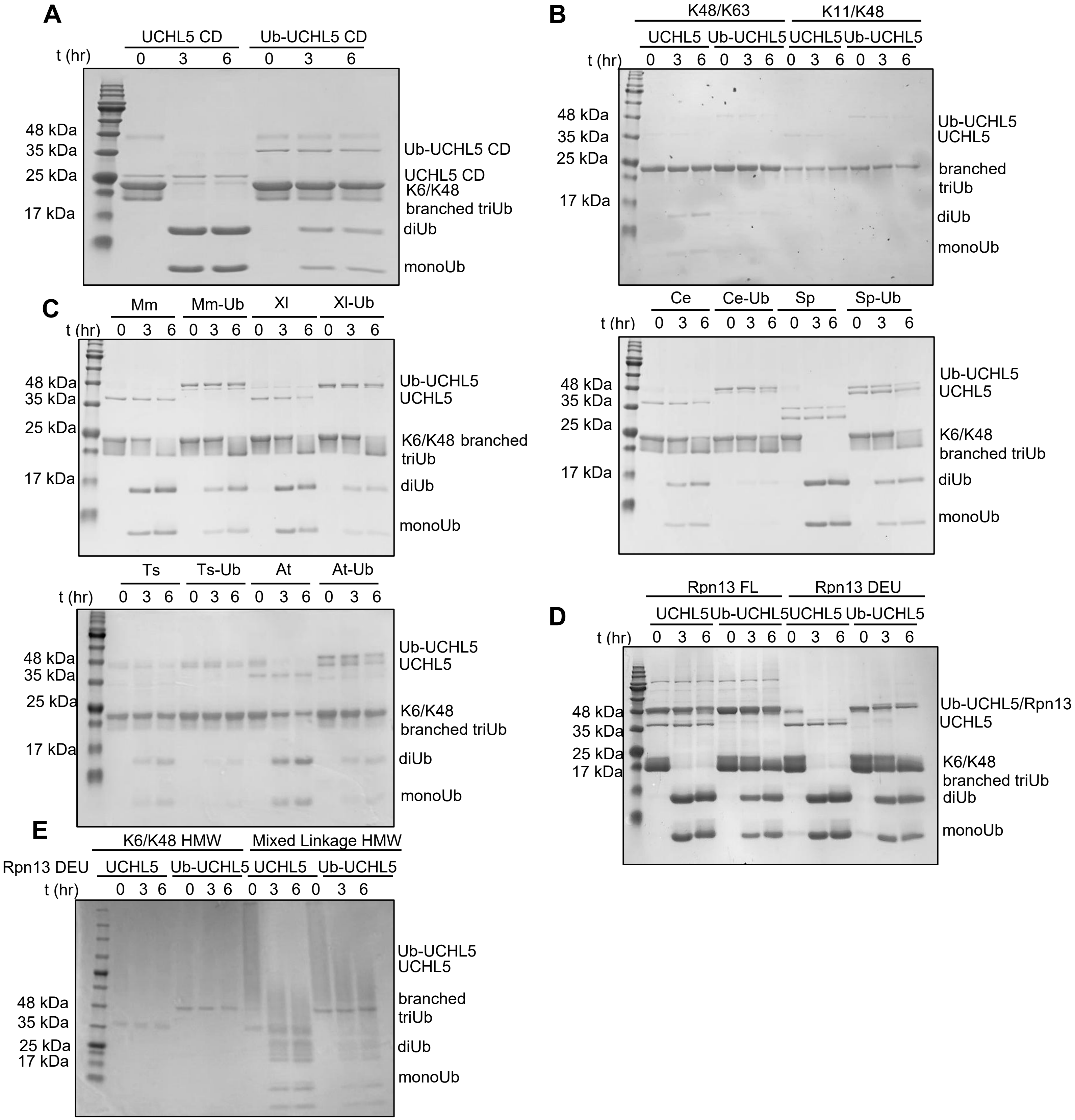

**Supplementary Figure 9.** SDS-PAGE analysis demonstrating (A) attenuated debranching activity of the N-terminally modified UCHL5 catalytic domain with Lys6/Lys48 branched Ub trimer substrate, (B) inhibition of debranching of Lys48/Lys63 branched trimer substrates by N-terminally modified UCHL5, (C) inhibition of chain debranching is largely conserved among N-terminally modified UCHL5 homologs relative to their unmodified counterparts, (D) association with Rpn13$^{DEU}$ and Rpn13 FL reinstates UCHL5 debranching activity toward Lys6/Lys48 branched triUb substrate, and (E) high molecular weight Ub chain substrates bearing Lys6/Ly48 and Lys6-, Lys11-, Lys63/Lys48 branch points.

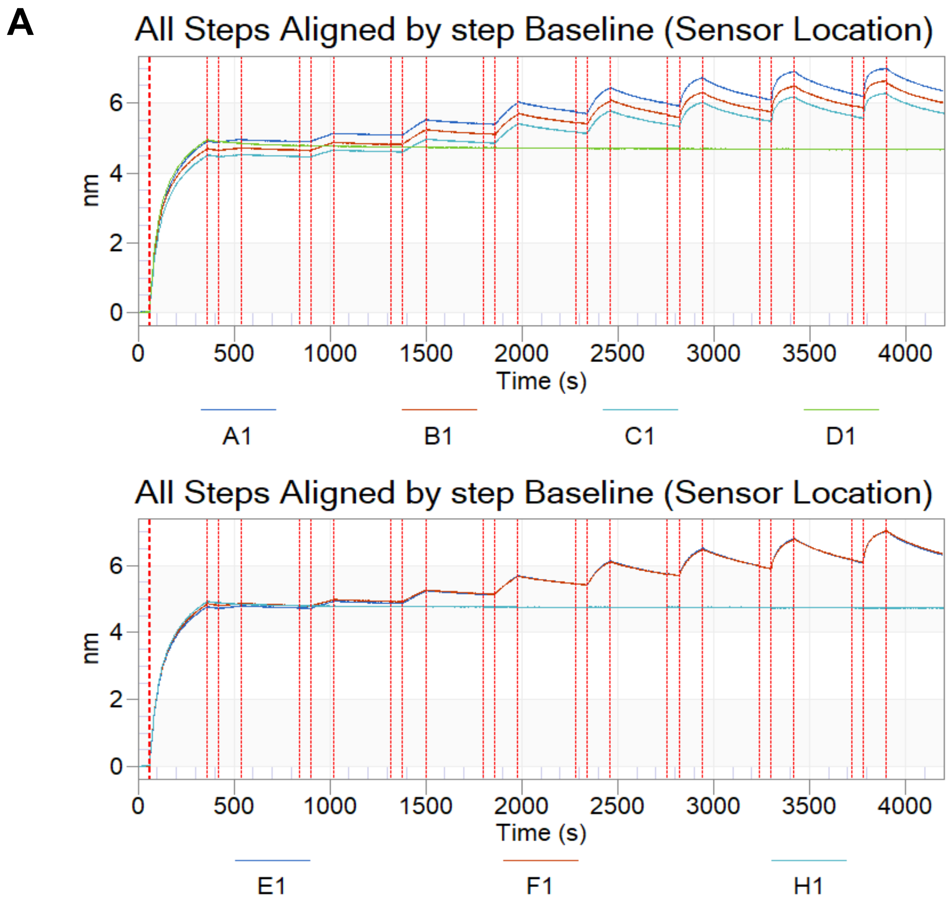

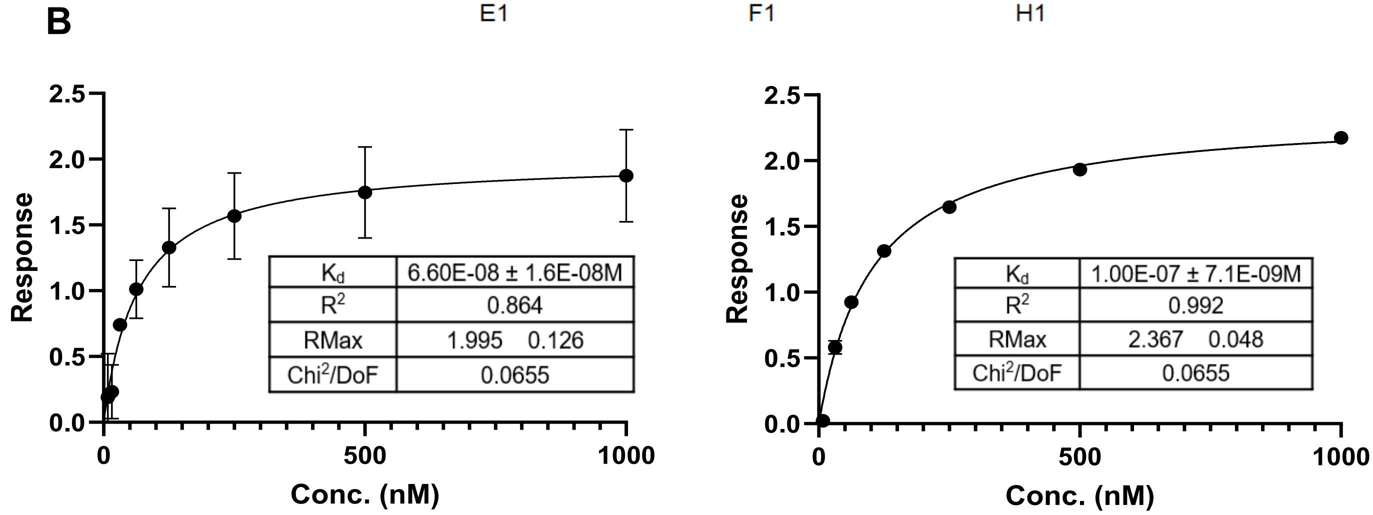

**Supplementary Figure 10.** A. Biolayer interferometry raw binding curves for UCHL5 (top) (n=3) and Ub-UCHL5 (bottom) (n=2) with Rpn13 FL B. Steady state kinetics with bait Rpn13$^{FL}$ indicate comparable dissociation constants for UCHL5 (left) and Ub-UCHL5 (right).

**A**

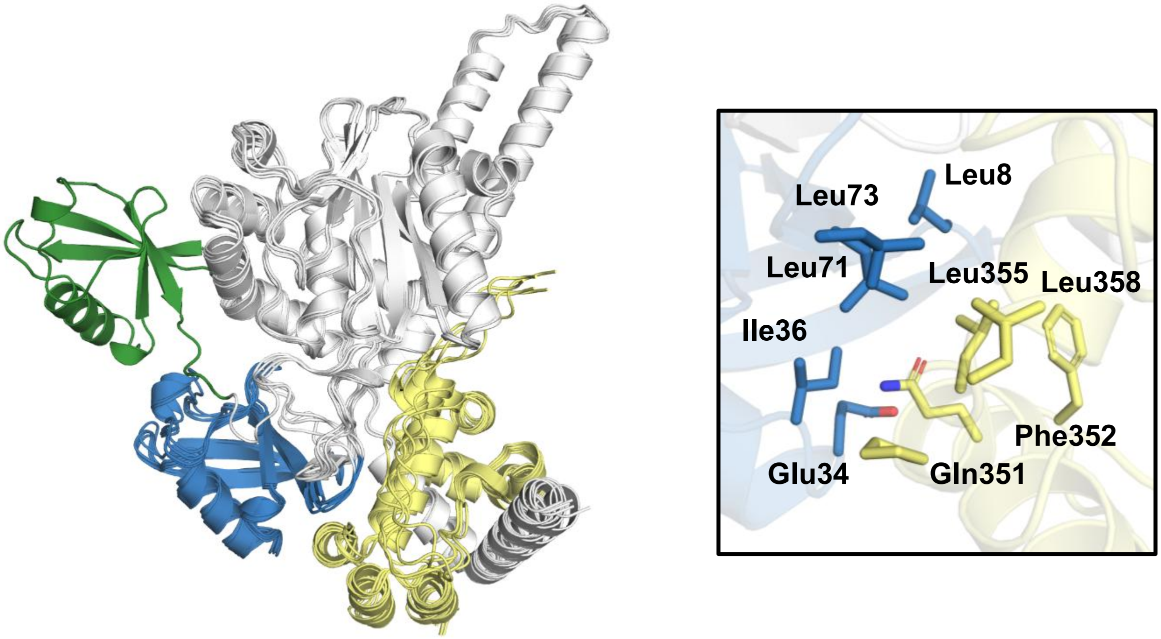

**B**

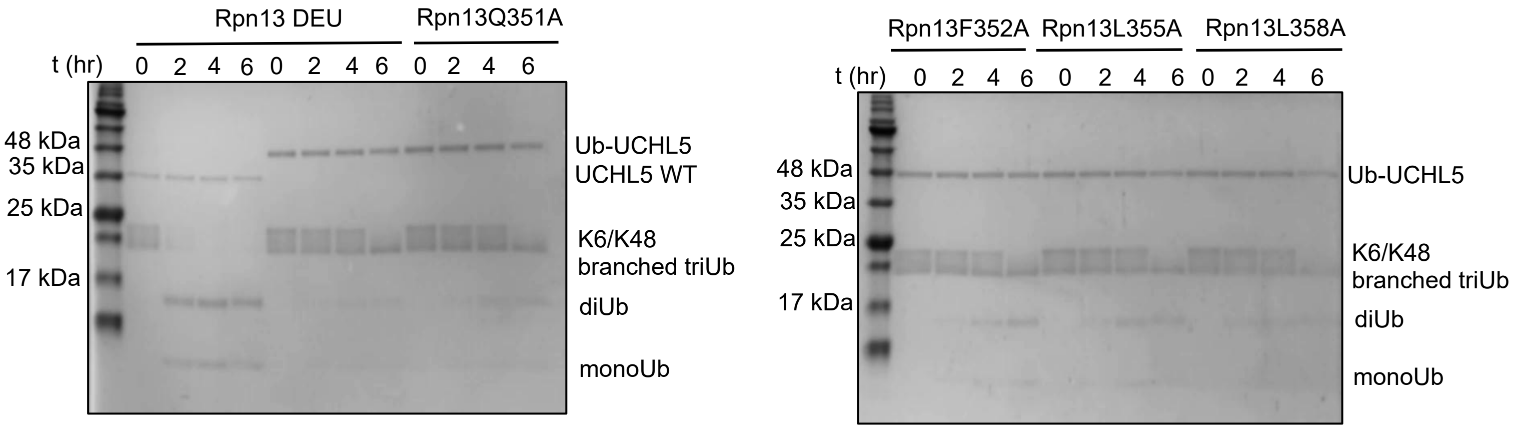

**Supplementary Figure 11.** A. Alphafold3 models of Ub-UCHL5 in complex with Rpn13$^{DEU}$ predict a shift of the N-terminal Ub from the back site to a neointerface generated by Rpn13 (left) Ub- Rpn13 interaction interface (right) B. Debranching activity of Ub-UCHL5 with Rpn13 mutants designed based on the predicted interface.

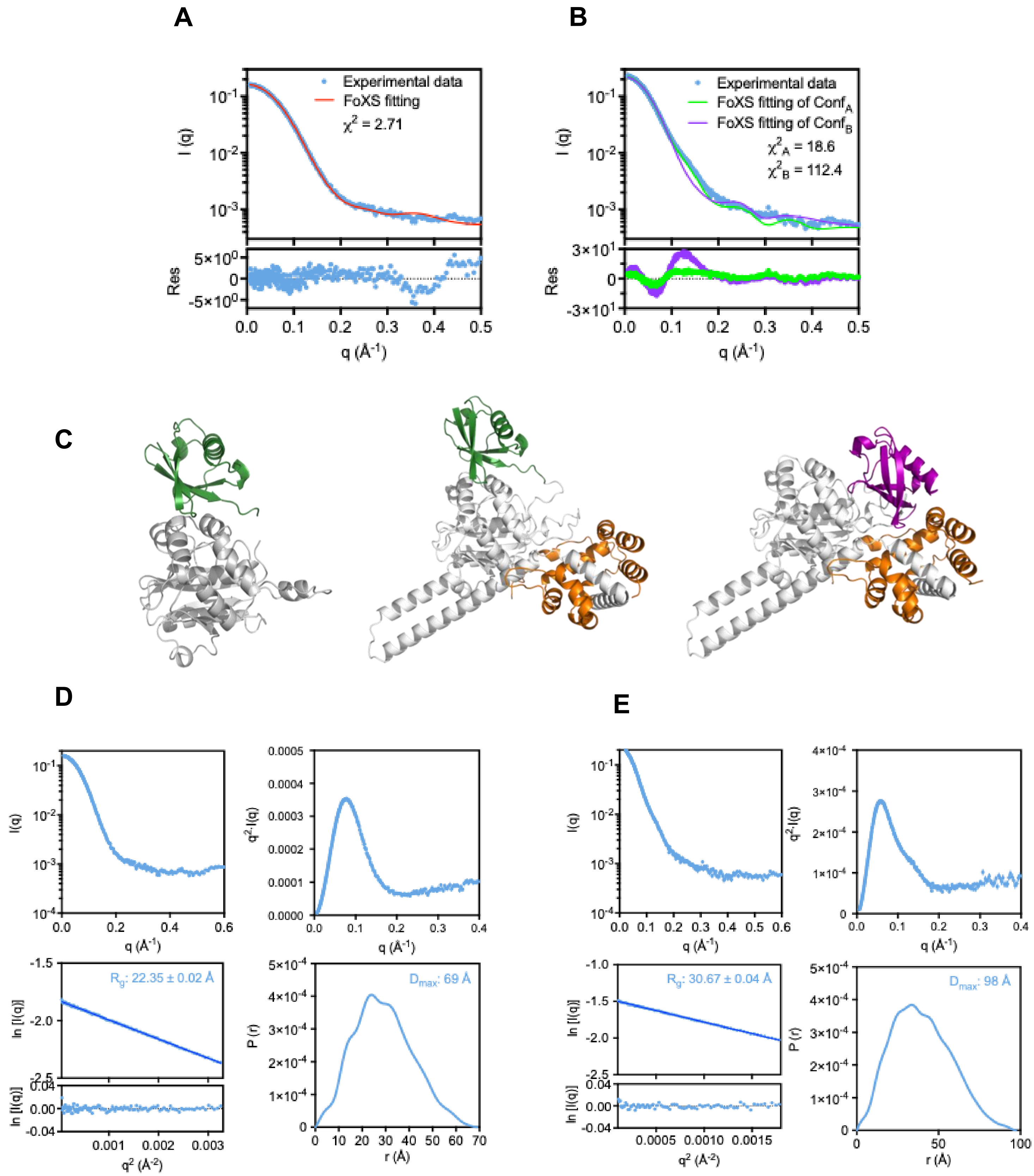

**Supplementary Figure 12.** SAXS analysis of N-terminally ubiquitinated UCHL5 variants. A. The experimental SAXS profile of Ub-UCHL5 CD is shown as blue dots. The theoretical SAXS profile generated by FoXS based on the atomic model of Ub-UCHL5 CD, modeled by Swiss-Model, is shown as a red line. The weighted residual is shown below. B. The experimental SAXS profile of Ub-UCHL5$^{FL}$ bound to RPN13$^{DEU}$ is shown as blue dots; the theoretical SAXS profiles derived from two distinct AlphaFold-predicted models generated by FoXS are shown as green and purple lines for conformation A ($conf_A$) and conformation B ($conf_B$), respectively. The weighted residuals are shown below with matching colors. C. Cartoon representation of the atomic model of Ub-UCHL5 CD, modeled by Swiss-Model (Left). UCHL5 CD is colored gray, and Ub is colored green. Cartoon representations of the AlphaFold model of Ub-UCHL5$^{FL}$ in complex with RPN13$^{DEU}$ in conf A (middle) and conf B (right). The two conformations differed by the relative positioning of Ub, colored in green and purple for conformations A and B, respectively. SAXS analysis of D. Ub-UCHL5$_{CD}$ and E. Ub-UCHL5$^{FL}$ in complex with Rpn13$^{DEU}$. SAXS profile (top left), Kratky plot (top right), Guinier plot with the fitting residuals shown below **(**bottom left)**,** and Pair-wise distance distributions, P(r), with the estimated maximum particle dimension, $D_{max}$, indicated above. (bottom right)
